# CK2 inhibitor, CX-4945, enhances BH3 priming and promotes apoptosis of venetoclax-resistant AML by targeting antiapoptotic proteins

**DOI:** 10.64898/2025.12.24.696284

**Authors:** Muhammad Daniyal, Rajesh Rajaiah, Upendarrao Golla, Marudhu Pandiyan Shanmugam, Koby Duke, Katherine Mercer, Yasin Uzun, Hannah Valensi, Jeremy Hengst, Sinisa Dovat, Yi Qiu, Suming Huang, Chandrika G Behura

## Abstract

Acute myeloid leukemia (AML), the most common hematologic malignancy, generally has a poor prognosis. Despite initial favorable responses to the BCL2 inhibitor venetoclax (VEN), remission is transient, and AML is eventually fatal. Resistance to VEN is primarily due to the overexpression of anti-apoptotic proteins, including MCL-1, BCL2L1 (BCL-XL), and BCL2A1. Casein kinase II (CK2) is a serine-threonine kinase and a known suppressor of apoptosis. We and others have reported that protein kinase CK2 activity is high in leukemic stem cells (LSCs) and associated with resistance to chemotherapy. We have shown that the selective CK2 inhibitor, CX-4945, suppresses BCL-XL and has a significant anti-tumor effect in AML preclinical models. CK2 expression and activity are high in venetoclax-resistant AML (VR-AML) cell lines. Genetic and pharmacological inhibition of CK2 significantly altered VR-AML gene signature, decreased MCL-1 protein level, increased BH3 priming and sensitized VR-AML cells to apoptosis. More importantly, CX-4945 selectively targeted LSCs (CD34+CD38−) and chemoresistant (CD123+CD47+) subpopulation in VR-AML. CX-4945 combined with VEN decreased leukemia burden and prolonged the survival of VR-AML cell line-derived and patient-derived xenografts compared to either drug alone. The combinatorial treatment was well tolerated in mice without additional myelosuppression or organ toxicity. CX-4945 (silmitasertib) is being tested in several early-phase clinical trials against adult and pediatric cancers. These preclinical results support the use of CX-4945 in combination with VEN to overcome resistance to apoptosis and re-sensitize VR-AML to chemotherapy.

**Graphical Abstract:** 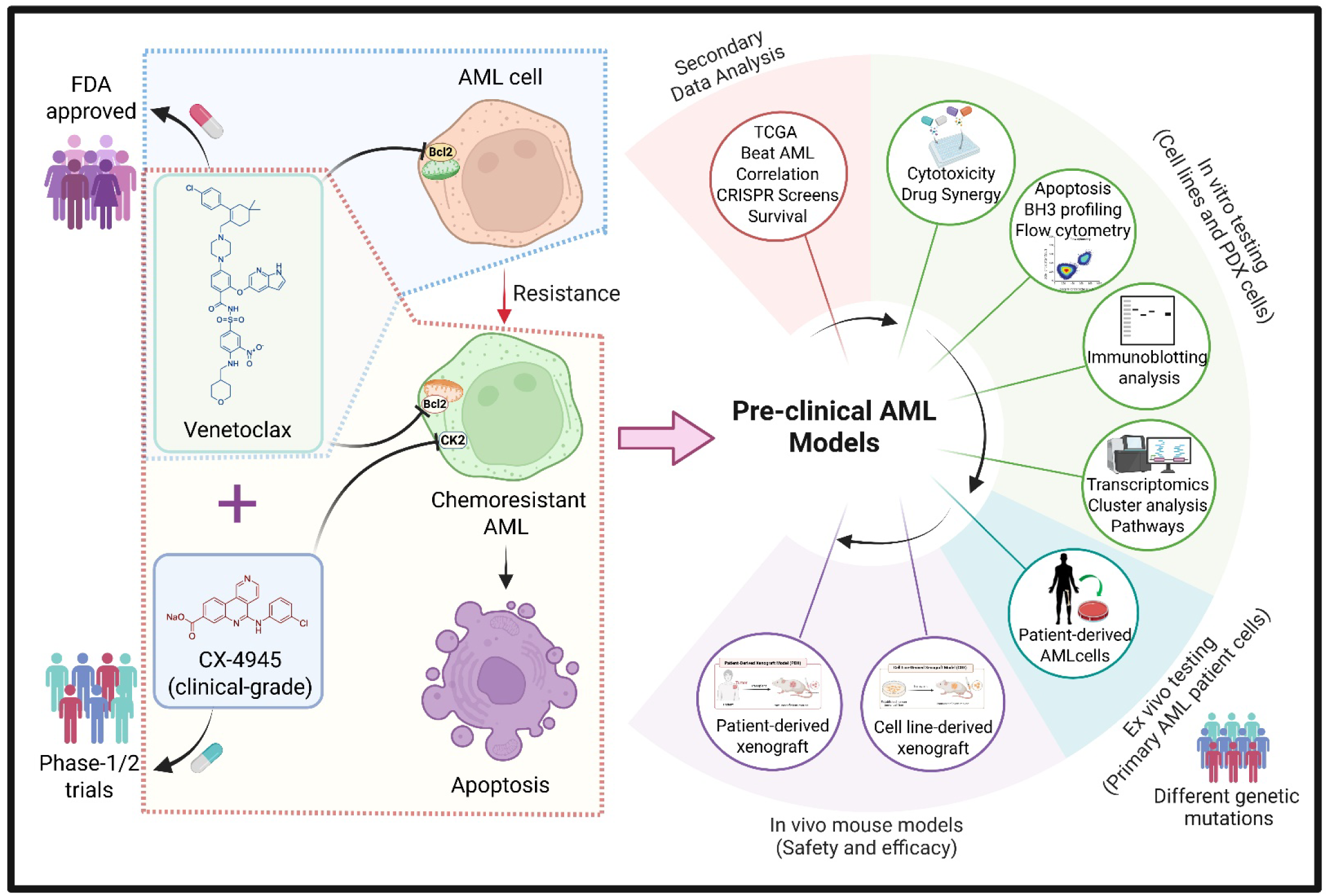

## Introduction

Acute myeloid leukemia (AML) is an aggressive hematopoietic malignancy with poor prognosis that results in less than 30% of adult AML patients surviving over five years [1]. It is heterogenous leukemia initiating from dysfunctional hematopoiesis with hindered proliferation and disrupted apoptosis [2, 3]. Cytotoxic chemotherapy and hematopoietic stem cell transplant remain the mainstay of treatment for AML [4, 5]. Venetoclax (ABT-199) is a small molecule that binds and antagonizes BCL-2 by mimicking the BH3 domain of pro-apoptotic proteins[6, 7]. Since AML cells exploit anti-apoptotic mechanisms for survival, the introduction of BH3 mimetic, ABT-199, a selective inhibitor of anti-apoptotic protein BCL2, successfully induced apoptosis in AML cells [8–10]. Venetoclax is FDA-approved for treating AML patients who are ineligible for intensive chemotherapy in combination with hypomethylating agents (HMAs) such as decitabine (DEC), azacytidine (5-AZA), or low-dose cytarabine (LDAC) [11–13]. Unfortunately, nearly half of the AML patients who received venetoclax-based therapy relapse within 18 months[14–16]. Therefore, sustaining durable remission with improved overall survival in AML still remains a challenge.

Adaptive resistance to venetoclax-based therapies in AML is multifactorial and involves mutations in BCL-2 resulting in loss of binding and altered survival dependence on BCL-2, or loss of function mutations in TP53, structural alterations in mitochondria, microenvironment changes in bone marrow, etc [17–21]. Apoptosis escape mechanisms mediated by increased expression of the BCL-2 family anti-apoptotic proteins MCL-1, and BCL-XL (encoded by gene *BCL2L1*) [18, 20–22] is a well-studied mode of venetoclax resistance and is actionable due to the availability of inhibitors of MCL-1. However, concomitant high expression of other anti-apoptotic proteins like BCL-XL and BCL2A1 renders AML cells resistant to MCL-1 inhibitors [22, 23]. Therefore, targeting MCL1 or BCL-XL alone may be inadequate to overcome venetoclax resistance [17, 19, 24]. Moreover, clinical use of direct BCL-XL or MCL-1 inhibitors is limited due to “on-target” dose-limiting toxicity of thrombocytopenia and cardiotoxicity respectively [7, 25]. Therefore, alternative ways of safely and effectively inhibiting anti-apoptotic and survival signaling pathways driving venetoclax resistance are needed. The existence of chemoresistant leukemic stem cells (LSCs) that are quiescent and unaffected by cytotoxic chemotherapy results in relapse [26]. Moreover, heterogeneity in the venetoclax resistance-driving mechanisms among AML genetic subtypes adds to the challenge of identifying a therapeutic target that is effective against most AML subtypes.

Protein Kinase/casein kinase (CK2) is a ubiquitous and constitutively active serine-threonine kinase, known to mediate cell survival and development [27–29]. An active CK2 enzyme is a tetrameric protein with two catalytic subunits (α or α’) and two regulatory subunits (β). Encoding genes *CSNK2A1*, *CSNK2A2,* and *CSNK2B* rarely show genetic alteration [30]. CK2 is highly conserved across species, and knockdown of *CSNK2A1* in mice is embryonically lethal, underscoring the essential function of CK2 in cell survival [31–33]. CK2 phosphorylates more than 300 substrates and regulates cell cycle, cell survival and apoptosis signaling pathways [27, 34, 35]. The elevated CK2 kinase activity has been implicated in the pathogenesis of several cancers, including hematological malignancies[36–38]. AKT, PTEN, NF-κB, STAT3, p53, IKAROS are among the major proteins directly or indirectly phosphorylated and regulated by CK2 [38–42]. Moreover, CK2 inhibition as an effective therapeutic strategy in lymphoid leukemia, lymphoma, multiple myeloma is proven in several preclinical studies[43–48]. In B cell leukemia, CK2 inhibition sensitizes B-cell leukemia cells to chemotherapy by dephosphorylating Ikaros transcription factor and potentiating its suppression of BCL-XL [49, 50]. In mantle cell lymphoma, CX-4945 overcomes venetoclax resistance via MCL-1 inhibition [45]. In AML, CK2 promotes leukemia stem cell survival via its regulation of AKT, NF-κB, STAT activation and IKAROS inactivation [51–53]. However, *CSNK2A1* deletion in the hematopoietic compartment does not significantly affect hematopoiesis in adult mice [54]. CK2α genetic (shRNA) or pharmacological (using CX-4945) inhibition showed *in vivo* anti-tumor activity in AML mouse models without significant myelosuppression [51, 53, 55]. Given the evidence of CK2 regulation of signaling pathways implicated in venetoclax resistance, including the antiapoptotic pathway, we hypothesize that CK2 inhibition will promote apoptosis and re-sensitize AML cells towards venetoclax.

CX-4945 (silmitasertib) is an orally bioavailable ATP-competitive, small-molecule inhibitor of CK2 with a favorable safety and tolerability profile in adult cancer patients [56]. Ongoing clinical trials are evaluating the safety and tolerability of silmitasertib in combination with chemotherapy in children and young adults with relapsed, refractory solid tumors (ClinicalTrials.gov NCT03904862, NCT06541262).

Here, we report the *in vivo* therapeutic efficacy of CX-4945 in venetoclax-resistant AML (VR-AML). CX-4945 treatment sensitized and primed VR-AML cells to BH3 peptides and increased cytochrome c release. CX4945, in combination with VEN, augmented apoptosis in VR-AML and primary patient samples. Moreover, the inhibition of CK2 reduced the expression of drug-resistant markers (CD47/CD123) and decreased leukemia stem cells (CD34+CD38−). CX-4945 treatment in combination with VEN prolonged the survival of mice xenografted with venetoclax-resistant AML and patient-derived xenograft cells. Our data thus provide a strong rationale with preclinical evidence for combinatorial blockade of both CK2 and BCL2 pathways in AML including VEN-resistant.

## Materials and Methods

### Drugs and Reagents

CX-4945 (#HY-50855B) and venetoclax (#HY-15531) were purchased from MedChemExpress (Monmouth Junction, NJ, USA). Recombinant human cytokines including SCF (#100-04), GM-CSF (#100-08), IL-3 (#100-80) were purchased from FUJIFILM Biosciences (Santa Ana, CA), while FLT-3 ligand (#PHC9411) and G-CSF (#PHC2031) were obtained from Thermo Fisher Scientific (Waltham, MA, USA). All other reagents, antibodies, and chemicals were of molecular biology grade and the details were provided in supplementary information.

### Cell culture

The human AML cell lines U937 (#CRL-1593.2), HL-60 (#CCL-240), THP-1 (#TIB-202), K562 (#CCL-243) were obtained from the American Type Culture Collection (ATCC) and MOLM-13 (#ACC554) cells were purchased from the German Collection of Microorganisms and Cell Cultures (DSMZ). The characteristics of AML cell lines were provided in **Table S1**. All AML cell lines were cultured in RPMI-1640 (#15-040, Corning, NY, USA) growth medium supplemented with 10% heat-inactivated fetal bovine serum (FBS) (#S11150; GeminiBio, West Sacramento, CA, USA) and 1X Penicillin-Streptomycin (#15140122; Gibco, Gaithersburg, MD) at 37°C in a humidified incubator with 5% CO_2_. Venetoclax-resistant (VR) derivatives of MOLM-13 and HL-60 cell lines were generated by continued subculture with increasing doses of VEN (starting from IC_50_) for 6-8 weeks. Cell lines were routinely confirmed negative for mycoplasma contamination and authenticated by short tandem repeat (STR) profiling (Promega, Madison, Wisconsin, USA). Each cell line was propagated for no more than 30 passages after initial thaw. The cell viability and count were measured by trypan blue exclusion assay using CellDrop automated cell counter (DeNovix, Wilmington, DE).

### Primary AML patient and Patient-derived xenograft (PDX) AML cells

De-identified and clinically annotated human AML patient samples derived from either bone marrow or blood were obtained from the Institutional Biorepository Core (RRID:SCR_025728) at the Penn State College of Medicine in compliance with institutional review board regulations. Cryo-preserved patient-derived xenograft (PDX) AML cells were obtained from collaborator and the director of Developmental therapeutic and pre-clinical core, Dr. Sinisa Dovat, who has obtained the samples from Cincinnati children’s hospital after MTA. All the primary and PDX AML cells were cultured in RPMI-1640 medium supplemented with 25% FBS, 1X penicillin/streptomycin, and 10 ng/mL of cytokines (IL-3, SCF, FLT-3, G-CSF, GM-CSF) at 37°C in a humidified incubator with 5% CO_2_ as described earlier [57]. The clinical characteristics of primary AML and PDX cells are summarized in **Tables S2 & S3**. The cell viability was measured via trypan blue exclusion assay right after thawing as well as during the culture period using CellDrop automated cell counter (DeNovix, Wilmington, DE).

### Cell viability assays

The AML cell lines or PDX AML cells were seeded in 96-well flat-bottom plates at 1×10^5^ cells/mL in 0.1 mL/well and treated with increasing concentrations of VEN (0.03-10 µM) and CX-4945 (2-6 µM) in combination for 48 h. Following the treatment, Cell Proliferation Reagent WST-1 solution (#05015944001; Sigma-Aldrich, St. Louis, MO) was added (1:10 final dilution) and incubated for 2-3 h. Absorbance was measured at 440 nm and 650 nm (reference wavelength) by SpectraMax i3x Multi-Mode Microplate Reader (RRID:SCR_026346; Molecular Devices, San Jose, CA). Following background subtraction, absorbances were presented as % viability relative to vehicle (DMSO) control.

### Drug synergy analysis

Dose-response matrices generated from percent cell viability data after drug combination treatment were used to compute synergy scores. The synergy between two drugs was analyzed using the SynergyFinder Plus (https://synergyfinder.org/) interactive webtool [58]. We used zero interaction potency (ZIP) reference model that calculates the expected effect of two drugs under the assumption that they do not potentiate each other [59]. ZIP scores greater than 10, between 10 and −10, and below −10 are considered synergistic, additive, and antagonistic, respectively.

### Lentiviral shRNA transduction

Lentiviral particles containing four unique shRNAs for human *CSNK2A1* (#TL320317V) and *CSNK2A2* (#TL320318V) were procured from OriGene Technologies (Rockville, MD, USA). Molm-13/VR AML cells were transduced in a 12-well plate with shRNA lentiviral particles for either *CSNK2A1* or *CSNK2A2* at 1-3 MOI (Multiplicity of Infection) using 8 µg/mL of polybrene (#TR-1003-G; Millipore Sigma, Rockville, MD) and spinoculation at 1600 rpm for 60 min at 32°C. The cells were sorted for GFP by flow cytometry under BSL-2 conditions using a BD FACSAria SORP (BD Biosciences, San Jose, CA) instrument in Penn State College of Medicine’s Flow Cytometry Core (RRID:SCR_021134) after 72 h post-transduction and screened for target gene knockdown by qRT-PCR and western blotting as described below. The shRNAs with highest knockdown efficiency were selected to achieve double knockdown of both *CSNK2A1* and *CSNK2A2* genes in the target AML cell lines. The cells transduced with scramble shRNA were used as a control for follow-up experiments including western blotting and apoptosis assay.

### RNA sequencing analysis

Total RNA was isolated from the AML cell lines (MOLM-13, MOLM-13/VR) and PDX (2016-7) cells after 24 h of treatment with DMSO (vehicle), CX-4945, VEN, and CX+VEN combo using TRIzol reagent (#15596018; Invitrogen, Carlsbad, CA, USA). The quality and integrity of RNA was assessed using Agilent 2100 bioanalyzer (Agilent Technologies, Palo Alto, CA, USA). Paired-end RNA sequencing was carried out using NovaSeq X Plus platform at Novogene (Sacramento, CA). The reads were processed, aligned to the human reference genome (hg38), and visualized as described earlier [60]. Differential expression analysis was conducted using edgeR [61], where a quasi-likelihood negative binomial generalized log-linear model was fitted to the data via the glmQLFTest function. Prior to modeling, raw count data were normalized using the trimmed mean of M-values (TMMs) method. The Benjamini–Hochberg (BH) procedure was applied for multiple testing correction, and genes with an adjusted p-value below 0.05 and fold change above 1.5 were considered as significantly differentially expressed. The principal component analysis (PCA) plots were generated using SRplot web server [62]. Hierarchical gene clustering (k-means) analysis with normalized counts data and functional enrichment analysis (MSigDB hallmark gene, GO-biological process) was performed using a web-based RNAseqChef platform [63]. Enrichment of gene signatures for diseases/drugs in differentially expressed genes was analyzed using Enrichr web server [64].

### Quantitative Real-Time PCR

Total RNA was extracted from AML cell lines (MOLM-13 and MOLM-13/VR) using TRIzol reagent and reverse-transcribed into cDNA using iScript cDNA Synthesis Kit (#1708891; Bio-Rad Laboratories, Hercules, CA). cDNA amplification was performed with PowerTrack SYBR Green Master Mix (#A46110; Thermo Fisher Scientific, Waltham, MA, USA). The thermal cycling program consisted of the following steps: denaturation at 94°C for 30 sec and annealing/extension for 30 sec at 60°C for a total of 40 cycles. All qPCR reactions were run in technical triplicates within the same run. *GAPDH* was used as an internal control gene for normalization and relative expression levels were determined by the 2^−ΔΔCt^ method [65]. The sequences for primers used in this study were listed in **Table S4**.

### BH3 profiling and dynamic BH3 profiling (DBP)

BH3 profiling involves determining sensitivity of AML cells to BH3 peptides without drug perturbation, whereas dynamic BH3 profiling (DBP) determines sensitivity after exposure to CX-4945. Intracellular BH3 profiling was performed as previously described in detail [66]. Exponentially growing AML cell lines (MOLM-13, MOLM-13/VR) were first stained with viability dye (1:1000), Zombie Aqua Fixable Viability Kit (#423102; BioLegend), washed with phosphate-buffered saline (PBS), and then resuspended in 50 μL of MEB buffer (150 mM mannitol, 150 mM KCl, 10 mM HEPES-KOH, 5 mM succinate, 20 μM EDTA, 20 μM EGTA, 0.1% protease free BSA, final pH 7.5). BH3 peptides (BIM, BID, PUMA) were prepared 2X the desired concentration in MEB buffer supplemented with 0.002% Digitonin. The cells were mixed with 50 μL of BH3 peptides (0.1 – 100 µM) or 50 µM alamethicin (ALA) and incubated for 90 min at room temperature and then fixed. Following neutralization, anti–Cytochrome C antibody (#612310; BioLegend) was added at a 1:2000 final dilution in staining buffer and incubated at 4°C overnight before acquisition with BD LSRFortessa (BD Biosciences, San Jose, CA) flow cytometer. ALA was used as positive control for cytochrome c (Cyt c) release.

For DBP, exponentially growing AML cell lines (MOLM-13, MOLM-13/VR) were treated with 10 µM of CX-4945 or equivalent DMSO (vehicle) for 16 h. After the treatment, cells were collected for subsequent BH3 profiling as described above and mixed with BH3 peptides (final concentration of 1µM for BIM, PUMA and 10µM of BID). Sensitivity to BH3 peptides was measured as percent cytochrome c release loss as determined by FACS and calculated using the following equation: [cytochrome c loss = 100 − (% of cells within cytochrome c retention gate)]. The readout of BH3 profiling and DBP is a change in mitochondrial priming defined as “% delta priming” (% delta priming = % cytochrome c loss [drug/peptide] − % cytochrome c loss [DMSO]). DMSO was used as a negative control for cytochrome c release, whereas a control without the cytochrome c antibody was used as a positive control for 100% cytochrome c release.

### Flow cytometry

To assess apoptosis in AML cells, we used PE Annexin V Apoptosis Detection Kit with 7-AAD (#640934; BioLegend, San Diego, CA, USA) per manufacturer’s protocol. Briefly, human AML cell lines and PDX cells were treated with vehicle (DMSO), CX-4945, VEN, and CX+VEN at indicated doses. After 24 h of treatment, cells were washed twice with ice-cold PBS and resuspended in 1X Annexin V binding buffer before adding 5uL of each PE-Annexin V and 7-AAD. The cells were incubated for 20 min at room temperature in the dark and then analyzed by BD FACSCalibur (BD Biosciences, San Jose, CA). The data were represented as a fold change in % apoptosis (Annexin V +Ve) cells relative to vehicle (DMSO) control.

For immunophenotyping, AML cells (cell lines, primary cells, and spleen or bone marrow cells harvested from xenograft mice) were collected (0.2-1 ×10^6^) by centrifugation and washed with ice-cold PBS before addition of viability dye (1:1000) from Zombie Aqua Fixable Viability Kit (#423102; BioLegend). After 20 min incubation in the dark at room temperature, cells were washed with FACS staining buffer (Dulbecco’s PBS with 3% FBS) prior to blocking mouse or human Fc receptor (BD Biosciences, San Jose, CA) for 10 min. The cells were then stained with an antibody cocktail for 30 min in the dark at 4°C and washed twice with FACS staining buffer before data acquisition using the BD LSRFortessa (BD Biosciences, San Jose, CA) flow cytometer. To analyze apoptosis in AML subpopulations, the cells stained with antibody cocktail were incubated with BV711-Annexin V (#563972; BD Biosciences) for 20 min at room temperature in the dark before data acquisition using the flow cytometer. The data analysis was performed using the FlowJo v10.10 Software (BD Life Sciences).

The frequencies of different cell types in the bone marrow and spleen samples were calculated by FACS analysis as described previously [60]. Absolute cell numbers were calculated based on the total cell numbers obtained by trypan blue exclusion assay using CellDrop automated cell counter (DeNovix, Wilmington, DE). The absolute number of cells in the populations of interest was calculated using the formula absolute cell number = (number of cells in gate/number of cells in live cells) × total cell number in organs (BM or spleen). All antibody information including clones and vendors is summarized in **Table S5**.

### Immunoblotting

The AML cell lines were seeded at a density of 4 × 10^5^ cells/mL in a 6-well plate and left untreated or treated with vehicle (DMSO), CX-4945, VEN, CX+VEN at indicated doses for 24 h. The cells were collected by centrifugation, washed with ice-cold PBS and then whole-cell lysates were prepared using 1X cell lysis buffer (#9803; Cell Signaling, Danvers, MA) supplemented with 1 mM of PMSF protease Inhibitor (#36978; Thermo Scientific) and phosphatase inhibitor cocktail (#P5726; Sigma-Aldrich). After quantification of protein concentration using BCA assay kit (#23225; Thermo Fisher Scientific), an equal amount of protein (10-20 μg) from each sample was denatured at 95°C for 5 min in reducing SDS loading buffer (#7722; Cell Signaling), subjected to gel electrophoresis using 4–20% mini-PROTEAN TGX precast protein gels (Bio-Rad Laboratories, Hercules, CA) and transferred to Immobilon-FL PVDF membrane (#IPFL00010; Millipore Sigma). Following blocking with 5% non-fat dry milk in TBST (Tris-buffered saline with 0.1% Tween-20), membranes were incubated with primary antibodies (listed in **Table S6**) overnight at 4°C. After three TBST washes of each 5 min, the blots were incubated with secondary antibodies either anti-rabbit (#7074, Cell Signaling, RRID:AB_2099233) or anti-mouse IgG–horseradish peroxidase (HRP) conjugates (#7076, Cell Signaling, RRID:AB_330924) for 45-60 min at room temperature and washed three times in TBST. SuperSignal West Femto Maximum Sensitivity Substrate (#34095, Thermo Scientific) was used as enhanced chemiluminescent (ECL) substrate and the signals were detected using Azure 500 imaging station (Azure Biosystems, Dublin, CA) or Bio-Rad ChemiDoc MP imaging system (Bio-Rad Laboratories, Hercules, CA). A representative blot from at least two or three independent experiments was presented.

### *In vivo* efficacy experiments

All mouse care and experiments were conducted in accordance with protocols approved by the Institutional Animal Care and Use Committee (IACUC). To test in vivo efficacy of CX-4945 and VEN combination, we used both cell line-derived xenograft (CDX) and patient-derived xenograft (PDX) mouse models. To establish CDX and PDX models, MOLM-13/VR (0.25×10^6^ cells/mouse) and U937 (1×10^4^ cells/mouse) AML cell lines and PDX 2016-7 (0.5×10^6^ cells/mouse) cells were injected via tail vein into 6-8-week-old NOD Cg-Rag1tm1Mom Il2rgtm1Wjl Tg(CMV-IL3,CSF2,KITLG)1Eav/J (NRG-S) mice (The Jackson Laboratory, Bar Harbor, ME; RRID:IMSR_JAX:024099). The mice (both sexes) were randomized into four experimental groups: Vehicle (control), CX-4945 (100 mg/kg, p.o., twice daily), VEN (12.5 mg/kg, p.o., once daily), CX+VEN. CX-4945 solution is freshly prepared in sodium phosphate buffer (pH=7.8) and passed through 0.22 µm filter before every dosing. VEN solution was prepared per manufacturer’s instructions by sequential addition of 5% DMSO, 40% PEG300, 5% Tween-80, and 50% saline. We included extra cohort of PDX mice for all the experimental groups to harvest blood and tissue (spleen and bone marrow) after two-weeks of treatment for analysis by flow cytometry and immunoblotting. The drug treatment was started a week after cell transplantation and continued until study termination. The mice were regularly monitored and complete blood count (CBC) from peripheral blood was performed every week till the end of the experiment by Hemavet 950FS analyzer (Drew Scientfic, Plantation, FL, USA). After the treatment period, the main study cohort mice were monitored for overall survival and subjected to Kaplan–Meier analysis.

### Statistical analysis

Statistical tests were performed using GraphPad Prism 10 software. Comparisons between two groups were performed by unpaired student’s t-test, while comparisons among three or more groups were evaluated by analysis of variance (ANOVA) test. Gehan-Breslow-Wilcoxon test was used to compute p-value for Kaplan-Meier survival analysis. All data were represented as the means ± SEM (standard error of the mean) or SD (standard deviation) from two or three independent experiments and a p-value less than 0.05 considered as statistically significant. Details of the post hoc tests if any used to calculate p-values are provided in the respective figure legends.

## Results

### High expression and activity of CK2 correlates with poor AML prognosis and resistance to venetoclax

AML patients expressing high level of *CSNK2A1* show decreased overall survival compared to low *CSNK2A1* levels in TCGA data (**Figure 1A**)[67]. Interestingly, higher *CSNK2A1* levels were significantly associated with *TP53*, *IDH1*, *WT1*, and *CEBPA* mutations in the BeatAML cohort (**Figure 1B**)[68]. Mutations in *TP53* specifically associated with resistance to VEN in BeatAML cohort (**Figure 1C**) [68], suggesting a potential role for *CSNK2A1* in aggressive disease phenotype and adverse treatment response. Accordingly, *CSNK2A1* levels exhibited positive correlation with *BCL2L1* (BCL-XL) levels that mediate resistance to VEN-based therapies (**Figure 1D**)[68]. In a kinase domain-focused CRISPR dropout screening, several AML cell lines (VEN-susceptible and inherently VEN-resistant) showed stronger dependency on genes encoding protein kinase CK2 and its interactome for survival (**Figure 1E**)[69]. Further to understand the role of CK2 in modulating VEN activity, we looked at genome-wide CRISPR dropout screens after VEN exposure. Interestingly, loss-of-function genes encoding essential CK2 catalytic subunit (*CSNK2A1*) and regulatory subunit (*CSNK2B*) depleted significantly in both Molm-13 and OCI-AML2 cell lines following 14 days of VEN treatment (**Figure 1F**)[18, 70], suggesting that overexpression of CK2 could confer VEN-resistance and inhibition of CK2 synergize with VEN activity. Based on synthetic lethal vulnerabilities of CK2 encoding genes with VEN, we were intrigued to study CK2 role in VEN resistance setting and use its clinical-grade selective inhibitor CX-4945 to achieve potentiated antileukemic activity in combination with VEN. Next, we tested a panel of AML cell lines and patient-derived xenograft (PDX) cells representing various molecular subtypes against VEN **(Figure S1A-B)**. PDX cells were derived from pediatric relapsed/refractory AML patients [71]. Details about the cell lines are provided in the supplemental material (**Tables S1 and S2).** All cell lines were resistant to VEN (IC_50_ >1 μM) except Molm-13, HL-60, THP1, and PDX 2016-1. We generated VEN-resistant Molm-13/VR and HL-60/VR cells by exposing parental Molm-13 and HL-60 cells to high concentrations of VEN (**Figure S1C**). As expected, both Molm-13/VR and HL-60/VR AML cells displayed resistance to VEN with IC_50_ more than 10 μM (**Figure 1G**), and slower cell proliferation kinetics (**Figure S1D**) compared to their parental counterpart. Interestingly, Molm-13/VR and HL-60V/R cells exhibited cross-resistance to decitabine, commonly used in combination with VEN, for AML frontline therapy (**Figure S1E**). Molm-13/VR cells exhibited upregulation of pro-survival BCL2 members (*MCL-1*, *BCL2L1*, *BCL2A1*) and downregulation of pro-apoptotic *BAK* expression without affecting genes encoding major CK2 subunits when compared to Molm-13 cells (**Figure 1H, S1F**).

**Figure 1:**
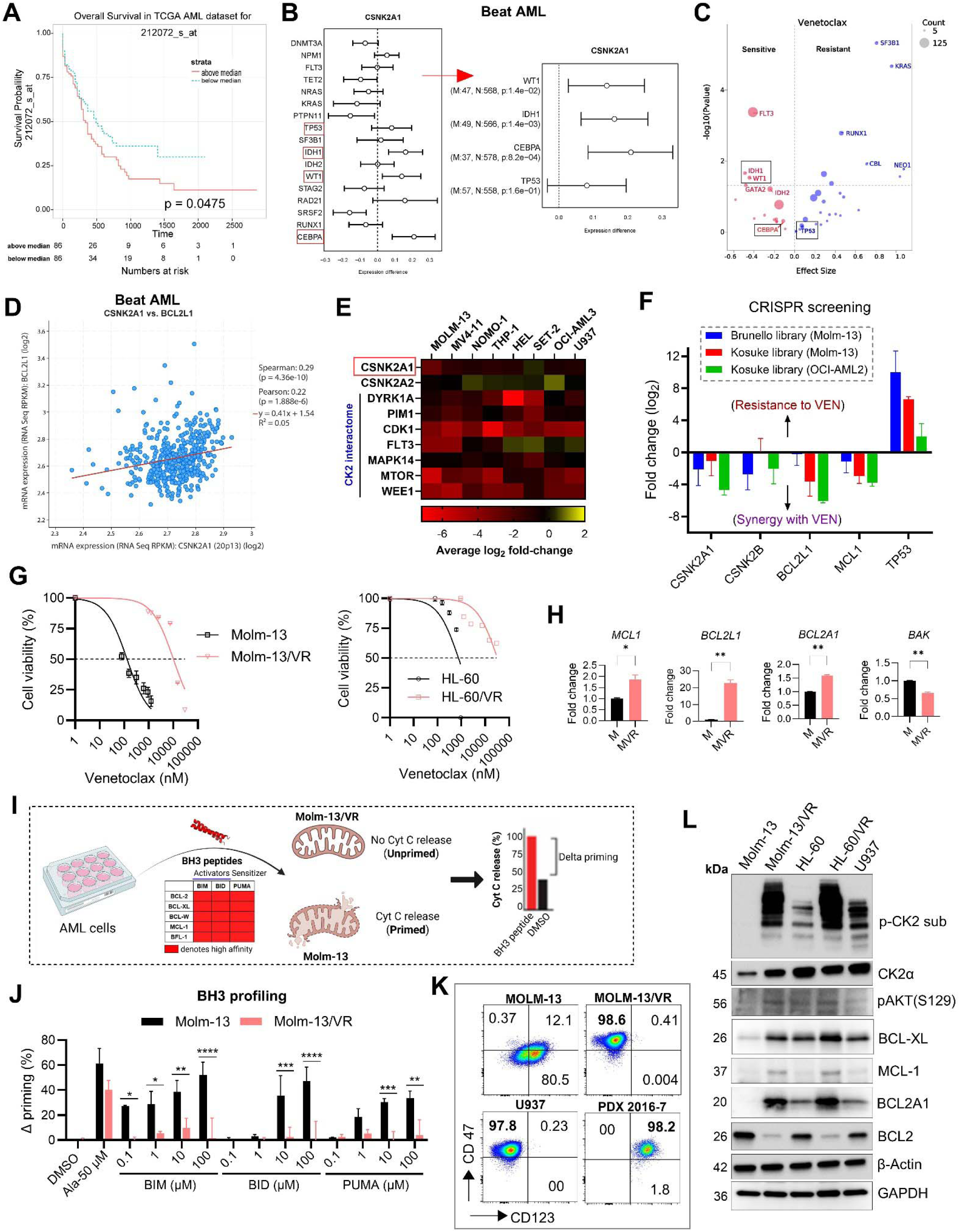
Characterization of CK2 expression and activity in AML and VEN resistance. **A)** Overall survival analysis of AML patients with low and high expression of *CSNK2A1* (probe ID: 212072_s_at) from TCGA data (AML vs normal; accessed from BloodSplot database at https://www.fobinf.com/). **B)** The plot depicts the median difference (95% confidence interval) of CK2α (*CSNK2A1*) expression in the presence or absence of mutations in the indicated gene calculated using Mann-Whitney tests. The gene mutations that significantly (p<0.05) associated with increased *CSNK2A1* expression were shown along with the sample number for mutated (denoted by M) and wildtype normal (denoted by N) in the BeatAML cohort. **C)** Volcano plot shows the association of gene mutation with venetoclax activity (accessed from BeatAML database at https://www.vizome.org/aml2/inhibitor/). Increased sensitivity is indicated by red, increased resistance indicated by blue as determined by the effect size (X-axis). **D)** Correlation between *CSNK2A1* (CK2α) and *BCL2L1* (BCL-XL) gene expression levels from Beat AML patient samples as determined by both Spearman and Pearson correlation coefficients. **E)** Heatmap summarizing the average log_2_ fold change of sgRNA abundance in kinase domain-focused CRISPR screening in AML cell lines (data adapted from [69]). **F)** Enrichment of individual sgRNAs for genes encoding CK2 catalytic subunit (*CSNK2A1*) and regulatory subunit (*CSNK2B*), BCL-XL (*BCL2L1*), MCL1, and TP53 plotted as log-fold change over control cells following 14 days of treatment with 0.5-1µM VEN in a CRISPR drop-out screening (data adopted from [18, 70]). Genes encoding BCL-XL, MCL1, and TP53 were used as known controls that alter VEN activity. Data are presented as Mean ± SEM (n=4-5 sgRNAs targeting each gene). **G**) Parental and VR-AML cell lines were treated with different concentrations of venetoclax for 48 h and assessed for cell viability using WST. **H)** The basal level expression of indicated genes was assessed in different AML cell lines by qRT-PCR. The data are presented as mean ± SEM (n=3 replicates from a representative run). *p<0.05 and **p<0.01 by unpaired t-test (Welch’s correction) denotes statistical significance. **I**) Schematic presentation of AML cells profiling with BH3 peptides (activators: BIM, BID; and sensitizer: PUMA) for the assessment of cytochrome c (Cyt C) release. Created with BioRender.com. **J)** Molm-13 and Molm-13/VR cells were tested for Cyt C release by priming with BH3 peptides and calculated the delta priming. The data are presented as mean ± SD (n=3) and analyzed by two-way ANOVA (Sidak’s multiple comparisons test). *p<0.05, **p<0.01, ***p<0.001, and ****p<0.0001 are considered statistically significant. **K)** The surface expression of chemoresistance markers (CD47 and CD123) on Molm-13, Molm-13VR, U937, and PDX 2016-7 cells was analyzed using flow cytometry. **L**) The basal level expression of CK2α, CK2 substrate phosphorylation, and other BCL2 family members in parental and VR-AML cell lines was analyzed by western blotting.

The differential gene expression analysis by RNA-sequencing showed significant downregulation of genes belonging to pro-apoptotic pathways (TNFα signaling, p53 pathway, apoptosis, STAT5 signaling, fatty acid metabolism) and upregulation of the genes belonging to heme metabolism that correlates with VEN resistance in Molm-13/VR cells compared to parental Molm-13 cells (**Figure S1G-H**). The differentially expressed genes (DEGs) and associated molecular gene signatures are common and comparable to Molm-13/VR transcriptome previously reported (GSE125403; **Figure S1H**)[72]. VEN resistance in Molm-13/VR cells was further confirmed by the level of cytochrome c (Cyt C) release in the presence of BH3 peptide priming (**Figure 1I**). BH3 peptides did not induce Cyt C release in Molm-13/VR compared to its parental Molm-13 cells (**Figure 1J**). Further, the surface expression of drug resistant markers was analyzed by flow cytometry. The higher level of surface expression of CD47 and CD123 in VR-AML cells has been reported [73]. The surface expression of drug-resistant markers was different in VR-AML cell lines and PDX cells. In fact, VR-AML cell lines exhibited high levels of CD47 compared to parental cell lines (**Figure 1K, S1I**). U937 cells, which are intrinsically resistant to VEN, also expressed high level of CD47 similar to Molm-13/VR cells (**Figure 1K**). Next, our immunoblotting analysis revealed that VR-AML cells have higher levels of CK2 subunits (α/α’/β) and CK2 activity as indicated by phospho-AKT (S129) and CK2-specific substrate phosphorylation (**Figure 1L, S1J**). As expected, VR-AML cells showed higher expression of pro-survival BCL2-family members (MCL1, BCL-XL and BCL2A1) with downregulation of BCL2 compared to parental AML cells. Altogether, our findings indicate that high levels and activity of anti-apoptotic protein kinase CK2 correlate with poor AML prognosis and VEN resistance, suggesting CK2 inhibition as a potential approach to inhibit AML progression, overcome resistance and enhance VEN activity.

### CK2 inhibitor CX-4945 in combination with VEN synergistically enhances apoptosis in VEN-resistant AML cells

Considering high CK2 activity in VR-AML cells and synthetic lethality with VEN, we reasoned to utilize CK2 selective clinical-grade inhibitor (CX-4945) to potentiate VEN-induced apoptosis and cytotoxicity in VEN-resistant AML cells. The cell viability of VEN susceptible and resistant cells in the presence or absence of CX-4945 was tested to evaluate the involvement of CK2 in drug resistance. CX-4945 as a single agent was not effective in reducing the cell viability in Molm-13, Molm-13VR, HL-60, HL-60VR and U937 cells (**Figure 2A**). Similarly, venetoclax as a single agent was also not effective in reducing cell viability of VEN-resistant cells tested. However, the CX-4945 (3 and 6 μM) combination with VEN significantly reduced the cell viability. Interestingly, the IC_50_ value of VEN in Molm-13/VR cells was significantly reduced to below 1 μM in the presence of CX-4945 (**Figure 2A**). Similarly, CX-4945 enhanced the cytotoxicity in AML relapsed/refractory PDX cells in combination with VEN and reduced its IC_50_ significantly (**Figure 2A**). The synergetic interaction between CX-4945 and VEN was established using zero interaction potency (ZIP) reference model with SynergyFinder webtool. The positive ZIP synergy scores suggested that CX-4945 in combination with VEN exhibited synergistic activity (score >10) in most AML cell lines and PDX cells tested except Molm-13 and U937 cell lines that showed additive activity (score between 0-10) (**Figure 2B, S2A-D**). The treatment with CX-4945 and VEN combo significantly enhanced apoptosis indicated by synergistic increase in annexin V levels in all the tested AML cell lines (both VEN susceptible and resistant) and PDX cells compared to single agents alone (**Figure 2C-D, S2E-F)**. Further, CX-4945 in combination with VEN reduced the expression of chemoresistance marker CD47 on VEN-resistant (both acquired and inherent) AML cell lines (**Figure 2E and S2G**). Additionally, exposure to CX-4945 enhanced priming effect of BH3 peptides and increased Cyt C release in Molm-13 and Molm-13/VR cells (**Figure 2F-G**). Next, we generated Molm-13/VR cells with genetic knockdown (shRNA-mediated) of CK2 catalytic subunits (α and α’) to test CK2 role in modulating VEN activity and resistance (**Figure S3A-B**). The Molm-13/VR AML cells with knockdown of CK2α/α’ exhibited partial increase in apoptotic activity of VEN compared to control cells (**Figure S3C**). Overall, these data indicated that CX-4945 potentiated VEN activity in combination to induce synergistic cytotoxic effect in both VEN-sensitive and resistant AML cells.

**Figure 2:**
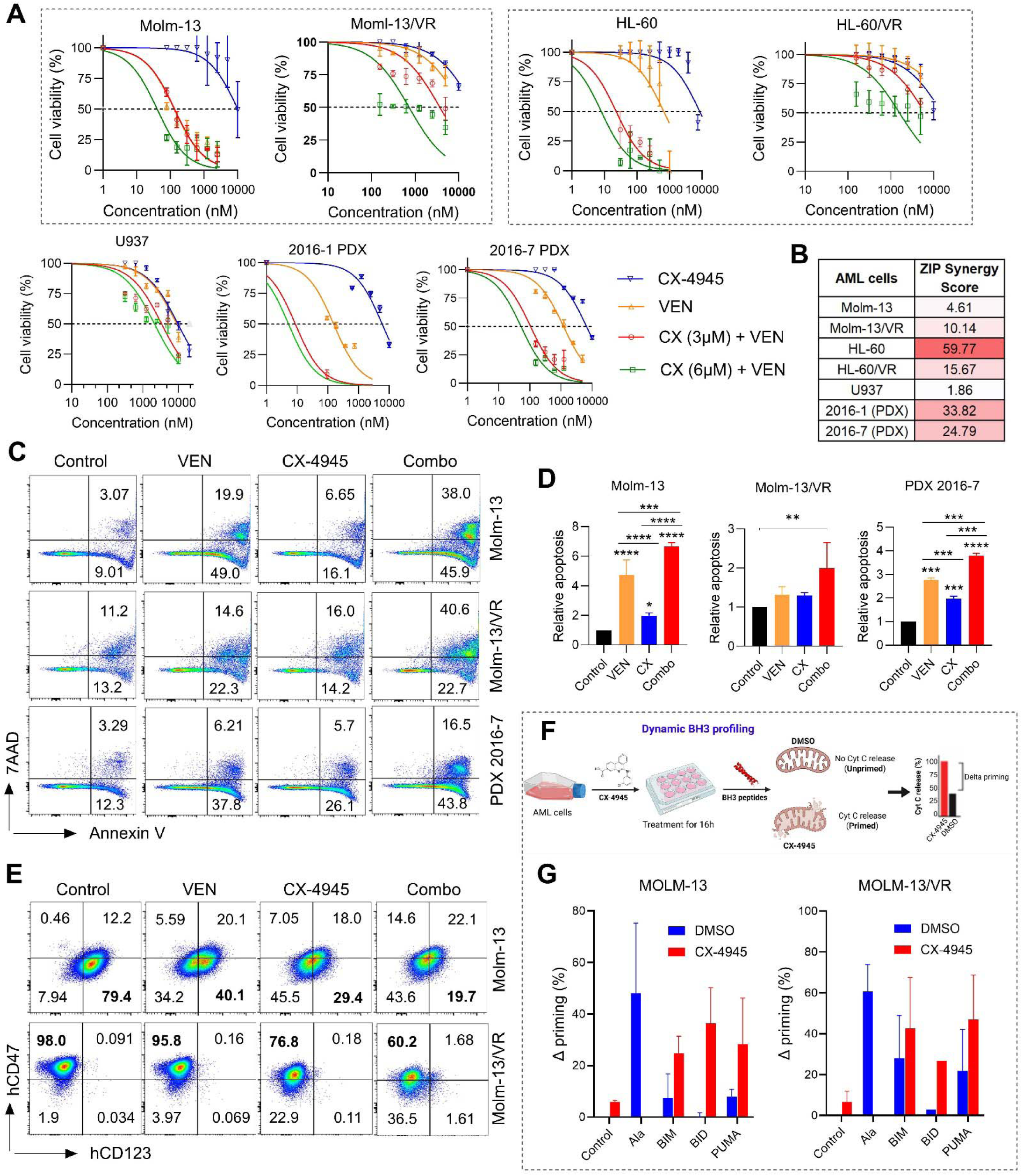
Synergistic cytotoxic activity of CX-4945 and VEN combination in AML cell lines and PDX cells. **A)** AML cell lines including both VEN-susceptible and resistant (Molm-13, Molm-13/VR, HL-60, HL-60/VR, U937), and AML PDX cells (2016-1, 2016-7) were treated with different concentrations of CX-4945 and VEN alone or in combination with indicated doses for 48 h. Cell viability was assessed using WST reagent and cell viability was calculated relative to vehicle-treated cells as 100%. **B)** ZIP synergy scores for CX-4945 and VEN combination in different AML cell lines and PDX cells were shown. ZIP scores greater than 10 indicates ‘synergy’, and 0-10 indicate ‘additive’ activity between CX-4945 and VEN. **C-D)** AML cells (Molm-13, Molm-13/VR, PDX 2016-7) were treated with CX-4945 and VEN alone or in combination for 24 h and stained with Annexin V/7AAD for analyzing apoptosis by flow cytometry (**C**). Relative apoptosis was calculated by normalizing to vehicle treated cells (**D**). The data are presented as mean ± SD (n=2-4). *p<0.05, **p<0.01, ***p<0.001, and ****p<0.0001 by one-way ANOVA (Holm-Sidak’s multiple comparisons test) denotes statistical significance. **E)** AML cell lines (Molm-13, Molm-13/VR) were treated with CX-4945 and VEN alone or in combination for 24 h and the surface expression of chemoresistance markers (CD47 and CD123) was analyzed by flow cytometry. **F)** Schematic showing the workflow for dynamic BH3 profiling to measure drug-induced priming of BH3 peptides in AML cells. The difference in Cyt c release (%) with and without drug treatment was presented as delta priming. Created with BioRender.com. **G)** Molm-13 and Molm-13/VR cells were tested for Cyt C release by priming with BH3 peptides with and without CX-4945 treatment and calculated the delta priming. The data are presented as mean ± SEM (n=2) and analyzed by two-way ANOVA (Sidak’s multiple comparisons test).

### The combination of CX-4945 and VEN targets MCL-1 and other anti-apoptotic BCL2-family members to induce apoptosis in VR-AML and PDX cells

MCL-1 (Myeloid CellLeukemia-1) is an anti-apoptotic BCL-2 family protein that plays critical role in hematopoiesis and blood malignancies. MCL-1 overexpression is a critical determinant for cross-resistance to standard therapies targeting BCL2/BCL-XL. The shorter isoforms of MCL-1 (S-short, ES-extra short) resulted by alternative RNA splicing or caspase-mediated cleavage inhibit MCL-1L and contribute to apoptosis in a BAX/BAK independent manner (**Figure 3A**). Since protein kinase CK2-mediated signaling through AKT/mTOR regulate MCL-1 levels at transcriptional and translational level, we performed immunoblotting analysis to check the levels of MCL-1 isoforms after CX+VEN combination treatment. Interestingly, our results showed that CX+VEN combo significantly reduced anti-apoptotic MCL-1L levels in Molm-13, Molm-13/VR, and PDX 2016-1 AML cells. Accordingly, the levels of pro-apoptotic MCL-1ES were increased in Molm-13, U937, and PDX 2016-7 AML cells (**Figure 3B**). Next, we asked whether MCL-1L levels are regulated at transcription or translation levels by Combo treatment. Quantification of MCL-1 transcripts after treatment showed lower levels in Molm-13 and PDX cells without significant changes in VR-AML cell lines (**Figure 3C**). These findings indicate that CX+VEN combo treatment regulate MCL-1 levels differently (low MCL-1L, high MCL-1ES) at both transcriptional and post-transcriptional levels to promote enhanced apoptosis in parental and VEN-resistant AML cells (**Figure 3B-C**). The effects of CX+VEN combo on MCL-1 regulation in AML cells are comparable with that of activity seen in multiple myeloma cells previously [45].

**Figure 3:**
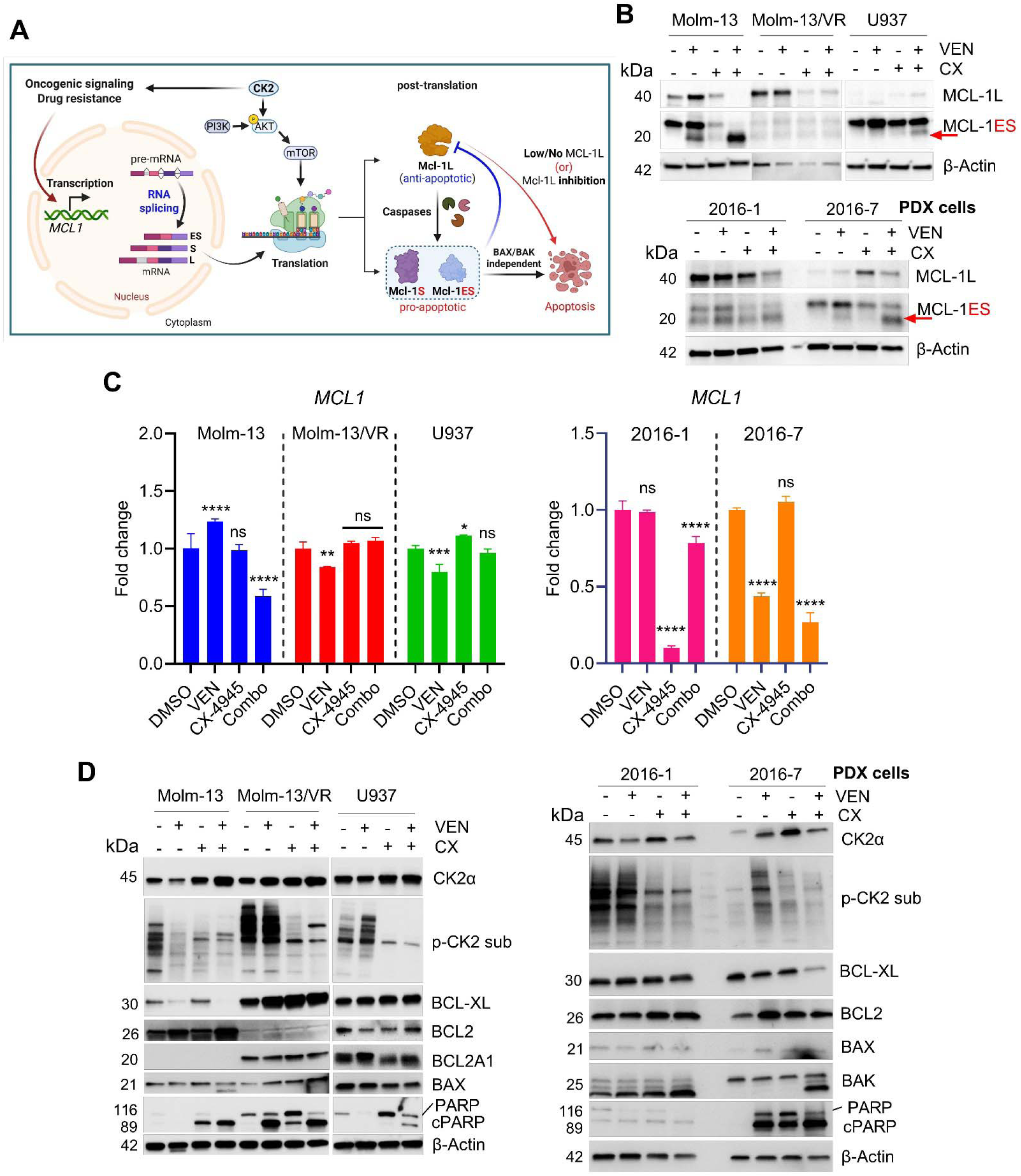
The effect of CX-4945 and VEN combination treatment on CK2 and BCL2-family member proteins in parental/VR-AML cell lines and PDX cells. **A)** The common mechanisms that govern cellular MCL-1 levels at transcription, translation, and post-translation levels are depicted. The signaling cascade mediated through CK2, PI3K/AKT, and mTOR regulate transcription and translation of MCL-1 isoforms (L-large, S-short, ES-extra short). MCL-1L (anti-apoptotic) undergoes caspase-mediated cleavage to form pro-apoptotic shorter MCL-1 isoforms (S & ES) that can induce cellular apoptosis independent of BAX and BAK. Created using BioRender.com. **B)** The levels of MCL-1 isoforms (L and ES) were measured by immunoblotting in AML cell lines and PDX cells after 24h of treatment with CX-4945 and VEN alone or in combination. **C)** Quantification of MCL-1 transcript levels after treatment with CX-4945 and VEN alone or in combination for 24h in indicated AML cell lines and PDX cells by qRT-PCR. The data are presented as mean ± SD (n=3) and analyzed by two-way ANOVA (Tukey’s multiple comparisons test). *p<0.05, **p<0.01, ***p<0.001, and ****p<0.0001 are considered statistically significant. **D)** Immunoblotting analysis of AML cell lines (Molm-13, Molm-13/VR, U937) and PDX (2016-1, 2016-7) cells after 24h of treatment with CX-4945 and VEN alone or in combination. β-Actin was used as a loading control. Representative blots from two to three independent experiments were shown.

In support of synergistic apoptosis induction with CX+VEN combo, we tested protein level of pro-survival and apoptotic members in different AML cell lines and PDX cells (**Figure 3D**). Interestingly, the level of cleaved PARP (substrate for caspases) was steadily increased with decreased CK2 activity (substrate phosphorylation) upon treatment with CX-4945 and VEN in all tested AML cell lines (Molm-13, Molm-13/VR, U937) and PDX cells. Moreover, the expression of anti-apoptotic BCL-XL was downregulated in Molm-13 and PDX 2016-7 cells (**Figure 3D**). The effect of CX+VEN combo treatment on other BCL2 members is not significant. Altogether, our results indicate that CX+VEN combo treatment decrease CK2 activity and other pro-survival members (MCL-1L and BCL-XL) to promote synergistic cytotoxicity in VEN-susceptible and resistant AML cells and PDX cells.

### CX-4945 treatment in combination with VEN differentially regulated apoptotic pathways in VEN-resistant and PDX AML cells

To assess transcriptional changes in VR-AML cells after BCL2 and CK2 co-inhibition, we performed RNA-Sequencing analysis in MOLM-13/VR (MVR) cells after 24 h treatment with CX, VEN, and Combo. The drug treatment resulted in significant transcriptional variability as indicated by principle component analysis of MVR cells (**Figure 4A**). We found 2731, 2327, and 3763 genes were differentially regulated at least 1.5-fold (adjusted p<0.05) by CX, VEN, and Combo treatment respectively **(Figure S4A**). Gene cluster analysis showed six distinct clusters (**Figure 4B**) enriched for different functional categories (**Figure 4C**). Specifically, cluster-4 is enriched for apoptosis, p53 pathway, cell cycle check point gene sets that were upregulated by Combo treatment. In contrast, gene sets related to mTOR signaling, adipogenesis, unfolded protein response (UPR), and hypoxia were enriched in clusters 5 and 6 with genes that were repressed by CX and Combo treatment (**Figure 4C**). Functional enrichment analysis of DEGs in MVR cells based on MSigDB hallmark gene set showed that pro-apoptotic gene signatures related to TNFα signaling, p53 pathway, apoptosis, E2F targets, G2M checkpoint, mitotic spindle formation were upregulated (downregulated in MVR cells) after CX and Combo treatment (**Figure 4D & 4E**). Interestingly, gene set related to heme metabolism that correlates with VEN resistance [74] was downregulated (upregulated in MVR cells) by Combo treatment. Moreover, gene sets related to pro-survival pathways including hypoxia, E2F/Myc targets, mTOR pathway, and glycolysis were downregulated by both CX and Combo treatment (**Figure 4D**). Next, we extended RNA-Seq analysis to PDX 2016-7 cells to understand mechanisms associated with potent antileukemic activity of CX and VEN combo. Interestingly, we found 912 and 1018 genes were differentially regulated at least 1.5-fold (adjusted p<0.05) by CX and Combo treatment respectively (**Figure S3A**). PDX 2016-7 cells failed to show significant transcriptional changes followed by VEN treatment. CX and Combo treatments found effective with significant changes in transcriptome (**Figure 4F** and **S4A**). Further cluster analysis of most-variable genes after treatment showed five distinct clusters (**Figure 4G**). The clusters 1 and 2 were enriched with gene sets related to oxidative phosphorylation, E2F targets, UPR, and mTOR pathway were downregulated by CX and Combo treatment (**Figure 4H**). Whereas, gene sets belonging to allograft rejection, antigen processing and presentation were upregulated by ex vivo treatment of PDX 2016-7 cells (**Figure S4B**). Functional enrichment analysis of DEGs showed that gene sets related to TNFα signaling via NFκB, p53 pathway, Interferon gamma and Interferon alpha response were upregulated by combo treatment. The gene signatures related to pro-survival mTORC1 signaling, STAT5 signaling, fatty acid metabolism, and glycolysis were significantly enriched in genes down-regulated by Combo treatment (**Figure 4I**). Moreover, the PDX 2016-7 cells showed downregulation of several genes that are highly expressed in VEN non-responder AML patients [75] along with upregulation of pro-apoptotic BCL2 family members and p53 targets after CX and Combo treatment (**Figure 4J**). As both MVR and PDX 2016-7 cells represent MLL-Rearranged (MLL-r) AML, we looked at relevant MLL-r gene signatures and found that CX and Combo treatment resulted in differential transcriptional regulation of genes belonging to cell morphology, defense response, cell migration, and immune response (**Figure S4C-F**). DrugSeq (Drug perturbation) enrichment analysis of DEGs from MVR and PDX cells revealed that transcriptional response with CX+VEN combo treatment resembled with that of CX-4945 (Silmitasertib), eIF4A3 inhibitor, PI3K/mTOR inhibitor, and multi-tyrosine kinase inhibitor (**Figure S4G-J**). Altogether, our global transcriptome analysis followed by functional overrepresentation analysis revealed several mechanistic insights including critical apoptotic and cell survival pathways that were targeted by CX as a single agent and CX+VEN combo to effectively overcome VEN resistance and induce cell death in both MVR (VEN acquired resistance) and PDX 2016-7 cells.

**Figure 4:**
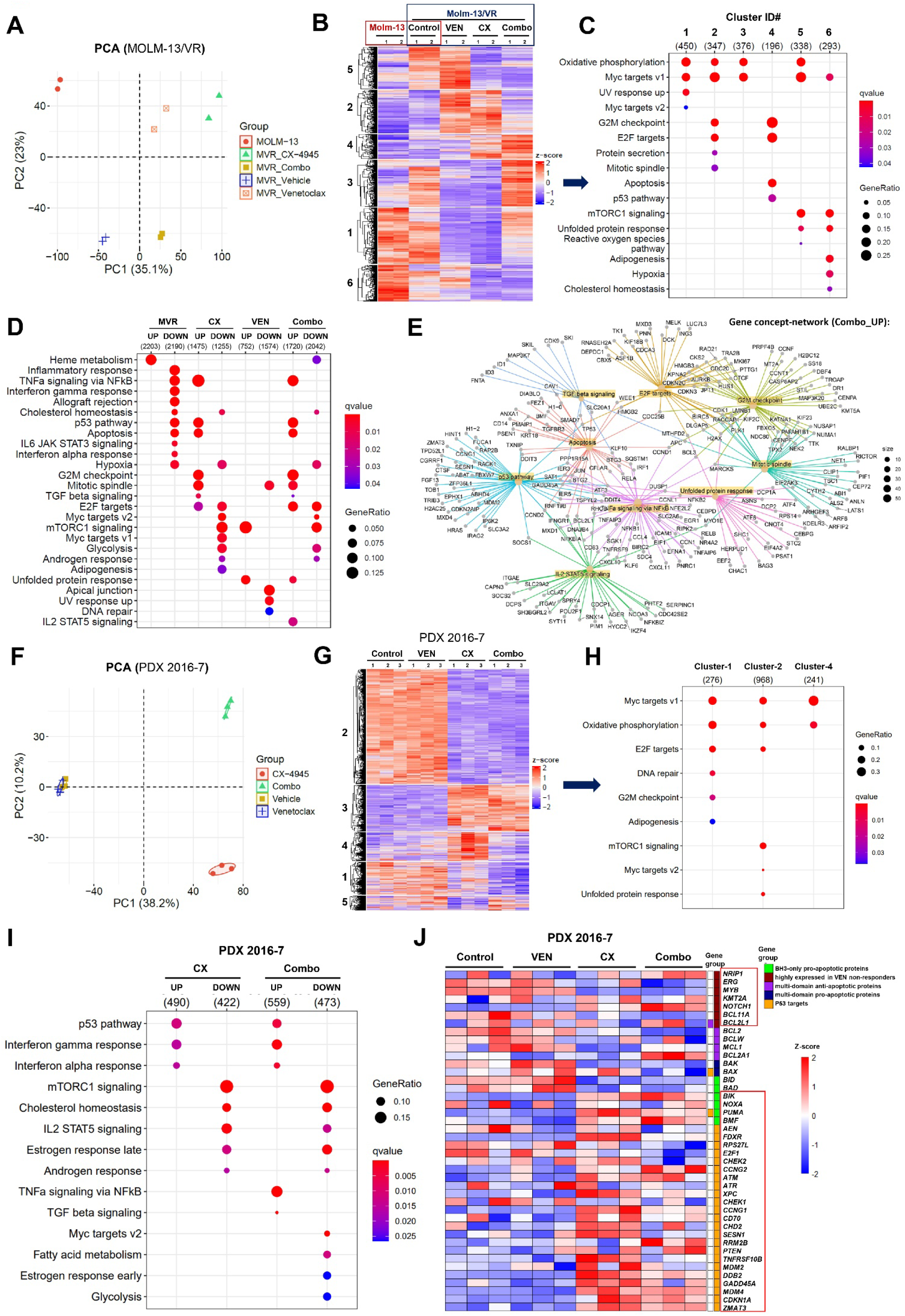
Differential gene expression and overrepresentation analysis with AML cells transcriptome. **A)** Principal component analysis (PCA) plot showing the distribution of Molm-13 (parental) versus Molm-13/VR (with and without drug treatment for 24h) AML cell lines (n=2 out of three replicates). **B)** Heatmap showing the hierarchical clustering (k-means) of most variable genes (n=2000) in Molm-13 and Molm-13/VR AML cell lines transcriptome. **C-D)** Dot plot displaying the top-ranked functional pathways (MSigDB hallmark gene set) overrepresented in different gene clusters obtained with k-means clustering as shown in ‘B’ (**C**) or the differentially expressed genes with fold change >1.5 and adj-p <0.05 (**D**). **E)** Gene-Concept network analysis depicting the gene-to-gene interconnection in the top-ranked functional pathways (MSigDB hallmark gene set) overrepresented with significant differentially expressed genes in Molm-13/VR cells with CX+VEN combo treatment (control vs. Combo). Node size refers to the number of genes in the enriched pathway. Genes shared between edges refer to terms belonging to multiple pathophysiological categories. **F)** Principal component analysis (PCA) plot showing the distribution of PDX 2016-7 (with and without drug treatment for 24h) AML cells (n=3). **G)** Heatmap showing the hierarchical clustering (k-means) of most variable genes (n=2000) in PDX 2016-7 AML cells transcriptome. **H-I)** Dot plot displaying the top-ranked functional pathways (MSigDB hallmark gene set) overrepresented in different gene clusters obtained with k-means clustering as shown in ‘G’ (**H**) or the differentially expressed genes with fold change >1.5 and adj-p <0.05 (**I**). **J)** Heatmap showing the expression profile of genes highly expressed in VEN non-responders (adapted from [75]), p53 target genes, and BCL2-family members (anti-apoptotic and pro-apoptotic) in PDX 2016-7 cells.

### Combination of CX-4945 and VEN is safe and synergistically prolongs the overall survival of AML CDX and PDX xenograft mice

To enhance the clinical translation relevance of our in vitro findings, we tested in vivo efficacy of CX-4945 and VEN combination in multiple pre-clinical mouse models generated with VEN-resistant (acquired or inherent) AML cells (Molm-13/VR, U937) and PDX 2016-7cells (**Figure 5A**). The immunocompromised NRG-S mice transplanted with PDX 2016-7 cells were treated with vehicle, CX-4945 (100mg/kg), VEN (12.5mg/kg), and CX+VEN combination to assess the safety and efficacy. Some animals from each experimental group were sacrificed after 2-weeks of treatment with single agents and CX+VEN, and assessed for spleen weight, CBC, engraftment of human CD45 and chemoresistant (CD47+CD123+) cell population in bone marrow (BM) and spleen (**Figure 5A**). Due to very high leukemia burden and short treatment period of 2-weeks, we did not see significant changes in the BM leukemia burden and chemoresisatant populations with combo treatment (**Figure S5A-D**). However, treatment with VEN and Combo effectively reduced AML PDX cells infiltration into spleen as shown by reduced spleen weight, percent hCD45 cells, leukemia burden, and chemoresisatant population (**Figure 5B-C & S5E-G**). The treatment with CX-4945 had very minimal to no effect on the overall CBC profile, while VEN and Combo treated mice displayed significantly lower WBC with improved RBC and platelet counts (**Figure 5D-F**). Moreover, immunoblotting analysis of spleen cells collected after 2-weeks of treatment showed that Combo treatment effectively reduced CK2 activity indicated by lower levels of AKT (Ser129) and NFkB p65 (Ser536) phosphorylation. Also, the levels of anti-apoptotic MCL-1L were greatly diminished with combo treatment compared to single agents (**Figure 5G**). The PDX mice treated with CX+VEN combo showed significantly (p<0.001) higher overall median survival (MS) of 41 days, relative to single agents (30-34 days) and vehicle control (27 days) groups (**Figure 5H**). Given the oral bioavailability and safety of clinical grade CX-4945 and VEN combination in PDX mice, we focused mostly on survival analysis with CDX mouse models described below.

**Figure 5:**
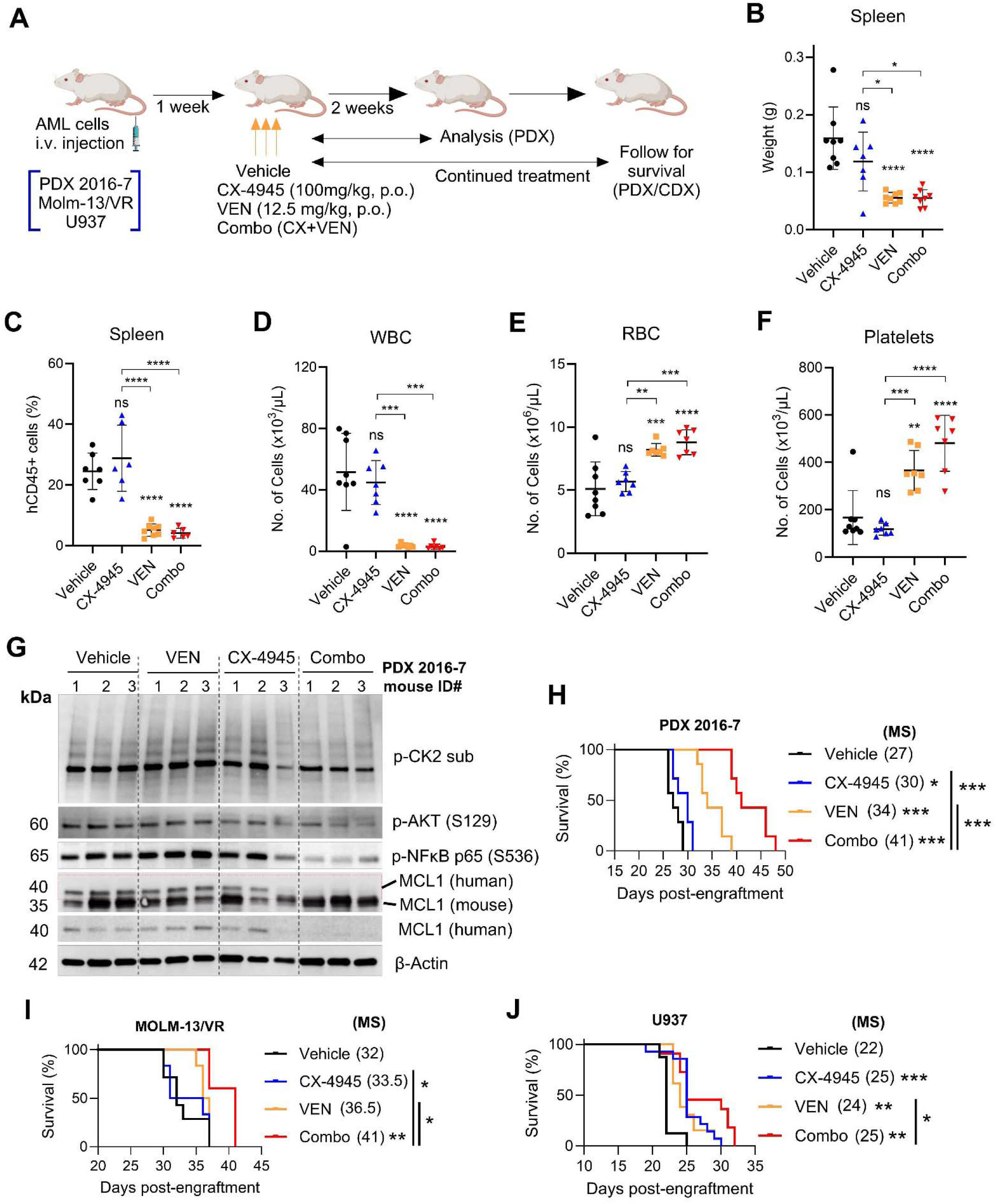
Anti-leukemic effect of CX-4945 in combination with venetoclax on AML-PDX, Molm-13VR and U937 cells xenograft in mice. **A)** Schematic depicting the treatment plan in AML patient-derived xenograft (PDX) mouse model. Similar scheme followed for cell line-derived xenograft (CDX) mice except the analysis after two-weeks of treatment. The NRG-S mice were irradiated and intravenously injected with AML cell lines Molm-13VR (0.25×10^6^ cells/mouse), U937 (1×10^4^ cells/mouse), and PDX 2016-7 (0.5×10^6^ cells/mouse) and randomized into experimental groups (Vehicle, CX-4945, VEN, and CX+VEN Como). The drug treatment started after one week of cell transplantation and followed up for survival analysis or analysis after two-weeks of treatment (only for PDX). **B-F)** After two weeks of drug treatment, mice injected with PDX 2016-7 cells were sacrificed and spleen weight was recorded (**B**). The cells collected from spleens were analyzed by flow cytometry to analyze percent hCD45 positive cells (**C**). **D-F)** CBC analysis was performed on PDX 2016-7 mice after two-weeks of drug treatment by Hemavet analyzer. The data for B-F are presented as mean ± SD (n=7-8) and analyzed by one-way ANOVA (Tukey’s multiple comparisons test). *p<0.05, **p<0.01, ***p<0.001, and ****p<0.0001 indicates statistical significance, while ‘ns’ denotes ‘not significant’. **G)** Immunoblotting analysis of spleen cells collected from PDX 2016-7 mice after two-weeks of treatment with CX-4945, VEN, and CX+VEN combo. The data for three different experimental animals was presented. **H-J)** Kaplan-Meyer survival analysis of 2016-7 PDX (**H**), Molm-13/VR CDX (**I**), and U937 CDX (**J**) mice after treatment with CX-4945, VEN and CX+VEN combo. The overall median survival (MS) days for each experimental group of PDX and CDX mice were provided next to the legend. *p<0.05, **p<0.01 and ***p<0.001 by Gehan-Breslow-Wilcoxon test indicates statistical significance.

Based on strong in vivo activity of CX+VEN combination in the PDX mice model, we were encouraged to test this combo in other CDX mouse models utilizing VEN-resistant AML cell lines including Molm-13/VR (acquired resistance model) and U937 (inherently VEN resistant). The CDX mice showed stable CBC profiles after 2-weeks of treatment with Combo regimen compared to vehicle control and single agents (**Figure S5H-I**). Intriguingly, next assessed the potency of the treatment of leukemic mice with CX-4945 in combination with venetoclax significantly enhanced the survival of mice injected with PDX-2016-7, Molm-13VR and U937 cells (**Figure 5F-H**). Interestingly, both the Molm-13/VR and U937 CDX mice showed significantly higher overall median survival after combo treatment compared to single agents and vehicle control groups (**Figure 5I-J**). All the CDX and PDX utilized in this study represent aggressive AML pre-clinical models with overall survival of 4-weeks following cell transplantation. Taken together, our in vivo efficacy studies highlight strong anti-leukemic activity and safety of CX+VEN combination in VEN-resistant preclinical AML CDX and PDX mouse models.

### CX-4945 treatment and in combination with VEN induced apoptosis in stem cells and drug-resistant primary AML patient cells

Primary AML cells were screened for the presence of leukemic stem cells (CD34+CD38−) and drug resistant markers (TIM3, CD47, CD123). Some of AML patient cells expressed stem cell marker, CD34 and positive for TIM3. On the other hand, all AML patient samples tested were either CD47+ or CD47+CD123+ (**Figure 6A and S6A**). Among tested patient samples, some are with FLT3, MLLr and IDH mutations and some are normocytic. AML cells with either Flt3 or MLL mutation exhibited increased CK2 substrate phosphorylation correlated with CK2a protein expression compared to normocytic AML patient cells and PBMCs from healthy donor (**Figure 6B**). Also, majority of tested AML patient cells showed higher levels of anti-apoptotic BCL-XL (**Figure 6B**). AML patient cells that exhibited high basal level apoptosis (∼>40%) without drug treatment were excluded from study (**Figure 6C**). AML cells with lower basal apoptotic activity were further assessed for cytotoxic effect of CX-4945 and VEN combination ex vivo. As a single agent, treatment with CX-4945 (2.5 and 5 μM) or VEN (0.25 and 0.5 μM) alone did not induce significant apoptosis in AML patient cells (**Figure 6D**). Notably, addition of CX-4945 significantly enhanced the apoptotic activity of VEN and AML patient cells exhibited higher apoptosis relative to control (**Figure 6D**). Interestingly, CX-4945 treatment alone was able to induce apoptosis in leukemic stem cell population (CD34+CD38−) without any effect of VEN (**Figure 6E**). However, CX-4945 alone did not induce significant apoptosis in drug-resistant AML cells (CD47+CD123+), but apoptosis was enhanced in combination with VEN in AML cells (**Figure 6E**). The combination was not effective to alter the frequency and induce apoptosis in other cell populations (**Figure S6B-C**). Together, these findings demonstrate promising cytotoxicity of CX-4945 and VEN combination in clinically-relevant primary AML patient cells *ex vivo*.

**Figure 6.**
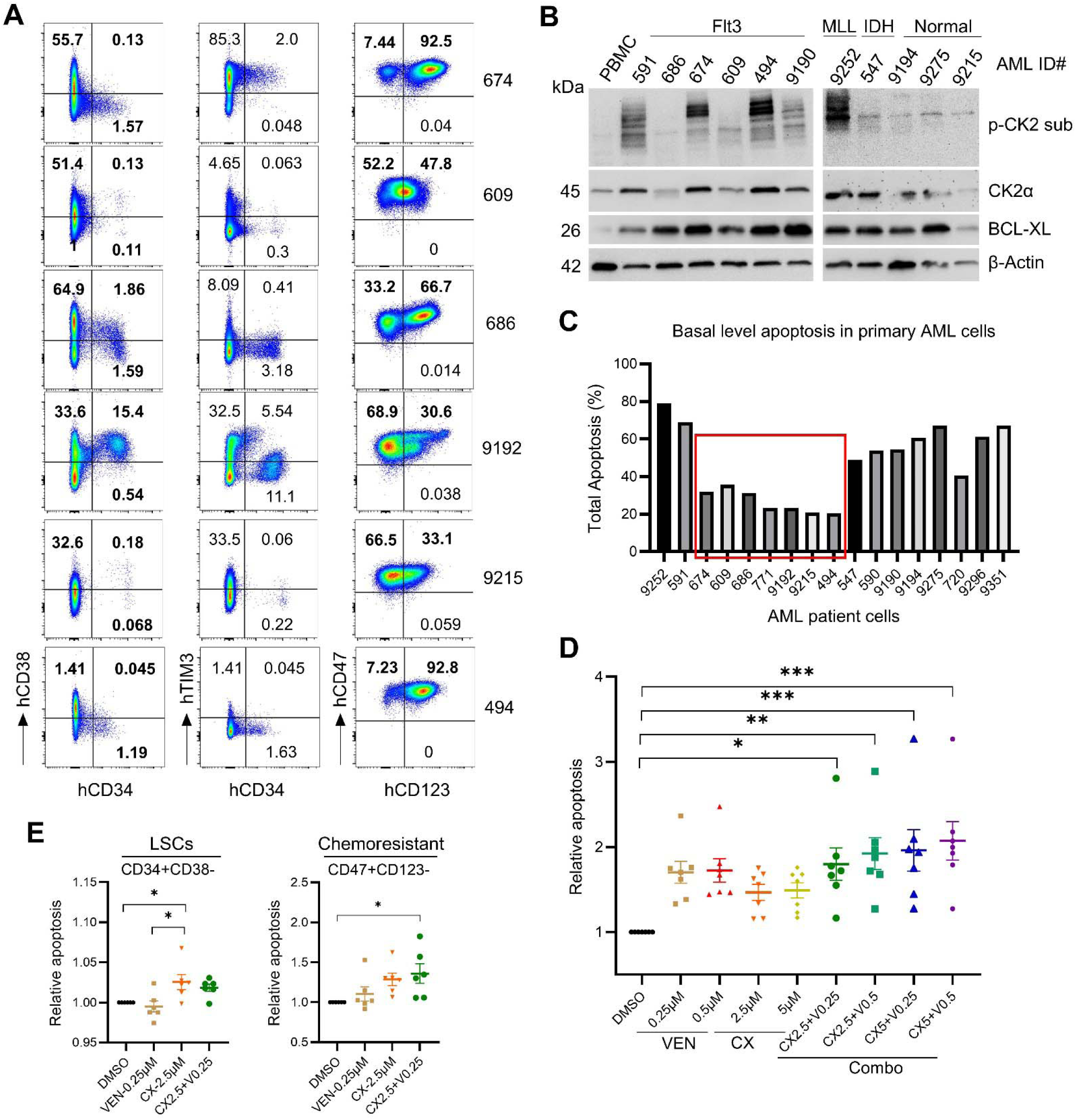
The effect of CX-4945 in combination with venetoclax on primary AML cells. **A)** AML cells from patients were surfaced stained with different fluorescent labeled antibodies and analyzed for the expression of markers for LSCs (CD34, CD38, TIM3) and chemoresistance (CD47 and CD123) by flow cytometry. **B)** The basal level expression of CK2α, BCL-XL, and CK2 activity (p-CK2 substrate) was measured in primary AML cells by immunoblotting analysis. **C)** The basal level apoptosis and viability of primary AML patient cells was assessed after 12-24h of post-thaw using Annexin V & dead cell kit for Muse cell analyzer. The patient cells with high basal level apoptosis were omitted and the ones used for further testing were highlighted using red square box. **D & E)** AML patient cells were treated with CX-4945 and VEN alone and in combination for 24 h and surface stained for different cell surface markers along with annexin V. The relative apoptosis in bulk cells (**D**), LSCs (CD34+CD38−) and chemoresistant (CD47+CD123-) subpopulations were calculated by normalizing the percent apoptosis in vehicle treated cells (**E**). The data are presented as mean ± SEM (n=6-7) and analyzed via one-way ANOVA (Tukey’s multiple comparisons test). *p<0.05, **p<0.01, and ***p<0.001 considered as statistically significant.

## Discussion

AML is generally treated by administering the combination of chemotherapeutic and targeted therapy drugs [76]. The treatment regimen with 7 days of cytarabine and 3 days of daunorubicin is considered the standard treatment practices [77–79]. A study from Papaemmanuil et al. established AML as a highly heterogeneous disease at the genetic-molecular level [80] and its therapy is increasingly shifting away from intensive chemotherapy towards personalized therapy [79]. The Food and Drug Administration approved several drugs that directly target mutations in FLT3, isocitrate dehydrogenase 1 and 2 leukemic drivers [81]. Other drugs including BCL-2 inhibitor, venetoclax that target cellular processes that are critical for the survival of AML cells [82]. The invent of treatment regimen with venetoclax in combination with hypomethylating agents is an outstanding development in AML treatment [8, 9]. However, the development to drug resistance knowingly impacts the effectiveness of venetoclax in relapsed or refractory AML [9, 83, 84]. Resistance to venetoclax in AML involve various genetic processes such as TP53 mutations, FLT3 kinase activation along with loss of function mutation in BAX [17, 18, 20, 85]. Apart from these genetic mutations, overexpression of anti-apoptotic proteins such as MCL-1, BCL-xL, and BCL2A1 during venetoclax resistance has been established [86].

Kinases play critical role in cellular processes to maintain homeostasis [87]. However, dysregulated kinases activity is responsible for survival of AML and venetoclax resistance [88]. Hence, kinases inhibitor in combination with standard drugs in use to treat AML have shown promising results in reversing drug resistance in AML. CK2 is one of the ubiquitous protein kinases involved in normal cellular processes [34]. Dysregulated CK2 activity has been reported in several pathological conditions including cancers [34]. Recently, the direct involvement of CK2 in the regulation of MCL1 in acute lymphoblastic leukemia and mantle cell lymphoma has been established [45, 89]. However, the direct involvement of CK2 in AML resistance to VEN has not been studied. In this study we reasoned and tested the possible involvement of CK2 in VEN resistance in AML.

The relationship between increased CK2 activity/expression and venetoclax resistance has been addressed by using a selective pharmacological inhibitor and genetic manipulation using CK2 specific shRNA. CK2 selective inhibitor, CX-4945, enhanced the cytotoxicity of VEN and reduced IC_50_ value, demonstrating the importance of CK2 in VEN resistance in AML cell lines and PDX cells. Moreover, the increased apoptosis and decreased surface expression of CD47/CD123 on VEN-resistant cells in the presence of CX-4945 further supported CK2-mediated drug resistance. As overexpression of anti-apoptotic proteins in VEN resistance, the priming effect of CX-4945 on BH3 mimetics and increased cytochrome c release clearly demonstrated CK2 mediation and drug resistance. The enhanced mitochondrial priming effect of anti-cancer drugs has been reported [83, 86]. In accordance with previous reports, the enhanced priming activity of CX-4945 supports consideration of CX-4945 as a combinatorial therapy for AML. Further, CX4945 in combination with VEN altered the gene signatures of several pathways including apoptosis, P53, mTORC1, cell cycle checkpoint, glycolysis, and E2F/Myc target genes analyzed using RNA-seq. In support of these findings, CX4945 treatment either alone or with VEN enhanced PARP cleavage and decreased anti-apoptotic MCL-1L levels, substantiating the advantage of CX4945 in treating venetoclax-resistant AML (**Figure 7**).

**Figure 7:**
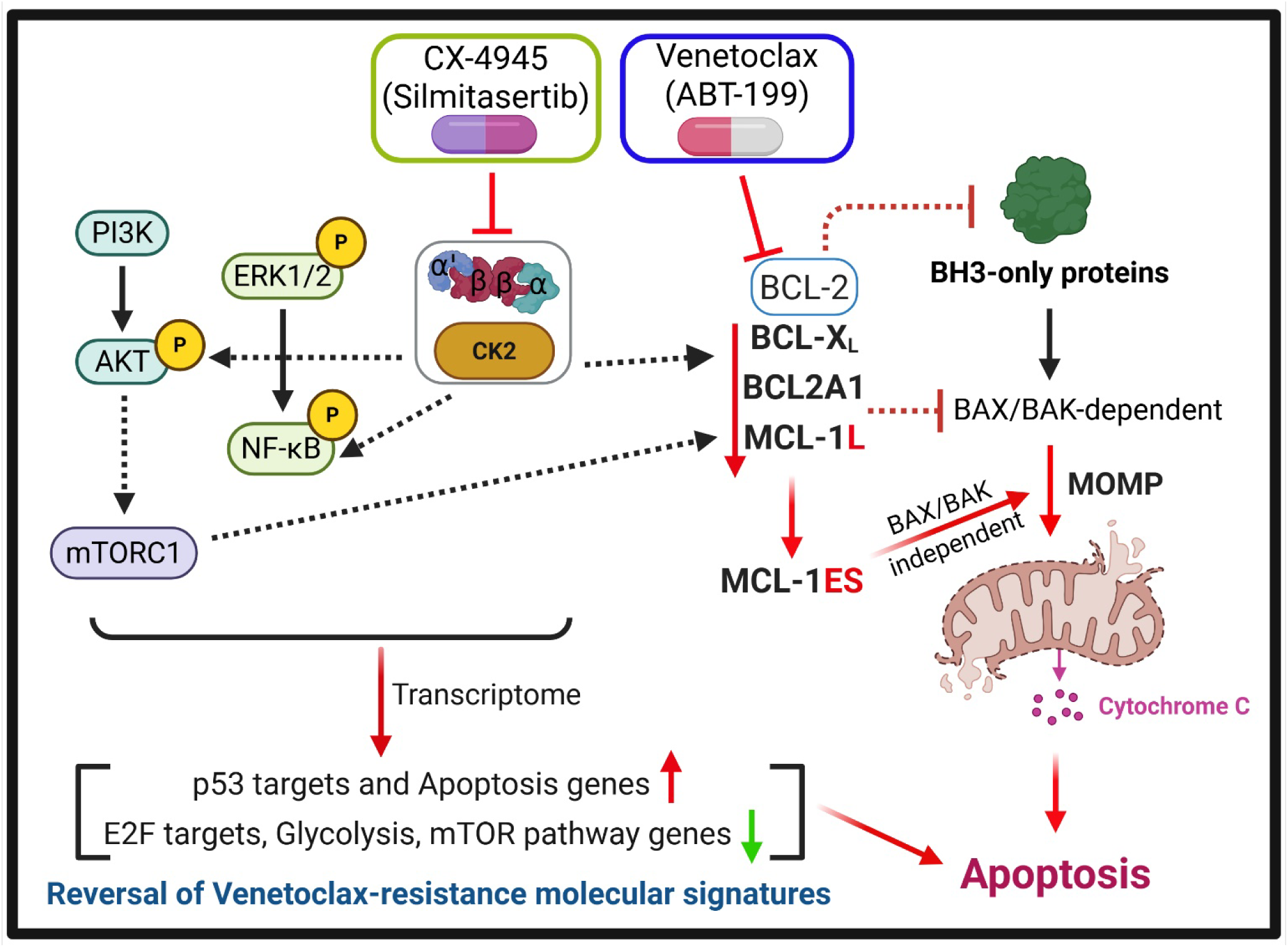
Summary model depicting the proposed mechanisms of action of CX-4945 and VEN combination in VR-AML cells. Inhibition of CK2-mediated signaling through CX-4945 negatively affect cell survival pathways (PI3K/AKT/mTOR, NF-κB pathways) and anti-apoptotic proteins (MCL-1L and BCL-XL). Downregulation of anti-apoptotic BCL2 members (MCL-1L, BCL-XL) further enhance BCL-2 dependency and enhance VEN-mediated apoptosis in VR-AML cells. Overexpression of pro-apoptotic MCL-1ES isoform that can mediate mitochondrial depolarization independent of BAX/BAK contribute to potentiate VEN activity. Moreover, the transcriptome of VR-AML cells followed by CX-4945 and VEN combination treatment showed reversal of molecular gene signatures associated with VEN resistance and result in potentiated apoptosis in VR-AML cells. The signaling targeted by CX-4945 and VEN are represented by dotted lines. Created using BioRender.com.

CX-4945 along with VEN enhanced the survival of mice xenografted with VR-AML cell lines (Molm-13/VR and U937) and AML PDX cells further proved the advantage of CX-4945 in treating AML as a combinatorial therapy. Several reports claimed that combining drugs targeting different pathways is beneficial with VEN or other standard therapies [90]. The common toxicities during the treatment of AML are anemia (reduced RBCs) and thrombocytopenia [91]. However, CX-4945 was able to specifically reduce the engraftment of AML cells confirmed by reduced CD47+CD123+ cells and maintain RBCs and platelets count. In fact, CX-4945 alone as well as in combination with VEN enhanced apoptosis of leukemic stem cell (CD34+CD38−) and drug resistant populations (CD47+/CD123+). The CD34+/CD38− immunophenotype specifies HSCs in AML that are quiescent in nature and is associated with poor prognosis in AML [92, 93].

## Conclusions

Innate and acquired resistance to BCL2 inhibitors in AML is associated with high expression and activity of CK2. BCL2i resistant AML is sensitive to CK2 depletion. Selective inhibitor of CK2, CX-4945, primes resistant cells to BCL2i induced apoptosis. We and others have shown anti-leukemic effect of CX-4945 in AML preclinical models. CX-4945 is currently being tested in Phase 1/2clinical trial for treatment of pediatric solid tumors. Here we show *in vivo* therapeutic efficacy of combining CX-4945 and venetoclax for treatment of resistant AML with specific targeting of chemo resistant and leukemia stem cells. These promising preclinical results support evaluation of this novel combination in clinical setting.

## Acknowledgments

Thanks to Developmental Therapeutics and the preclinical core and its faculty, Dr. Arati Sharma, and Dr. Diwakar B. Tukaramrao, for providing AML PDX cells. Penn State IPM team for sharing de-identified AML patient samples.

## Institutional Review Board Statement

All the animal experiments were conducted in accordance with guidelines and following protocols approved by the Institutional Animal Care and Use Committee at Penn State Hershey, Hershey, PA (IACUC # PROTO201901034). De-identified patient samples were provided by Institute of personalized medicine, at Penn State University College of Medicine (Hershey, PA, USA), and used in compliance with Institutional Review Board regulations.

## Informed Consent Statement

Not applicable.

## Data Availability Statement

The datasets analyzed and novel reagents used during the current study are available from the corresponding author upon request and after a material transfer agreement.

