## Supplementary data for "CK2 inhibitor, CX-4945, enhances BH3 priming and promotes apoptosis of venetoclax-resistant AML by targeting antiapoptotic proteins"

**Running title:** CK2 inhibition overcomes BCL2i resistance

^#^These authors contributed equally

^a^Present address: Eurofins Lancaster Laboratories, Lancaster, PA 17601.

^b^Present address: Maynard Children's Hospital at ECU Health Medical Center, Greenville, NC 27834.

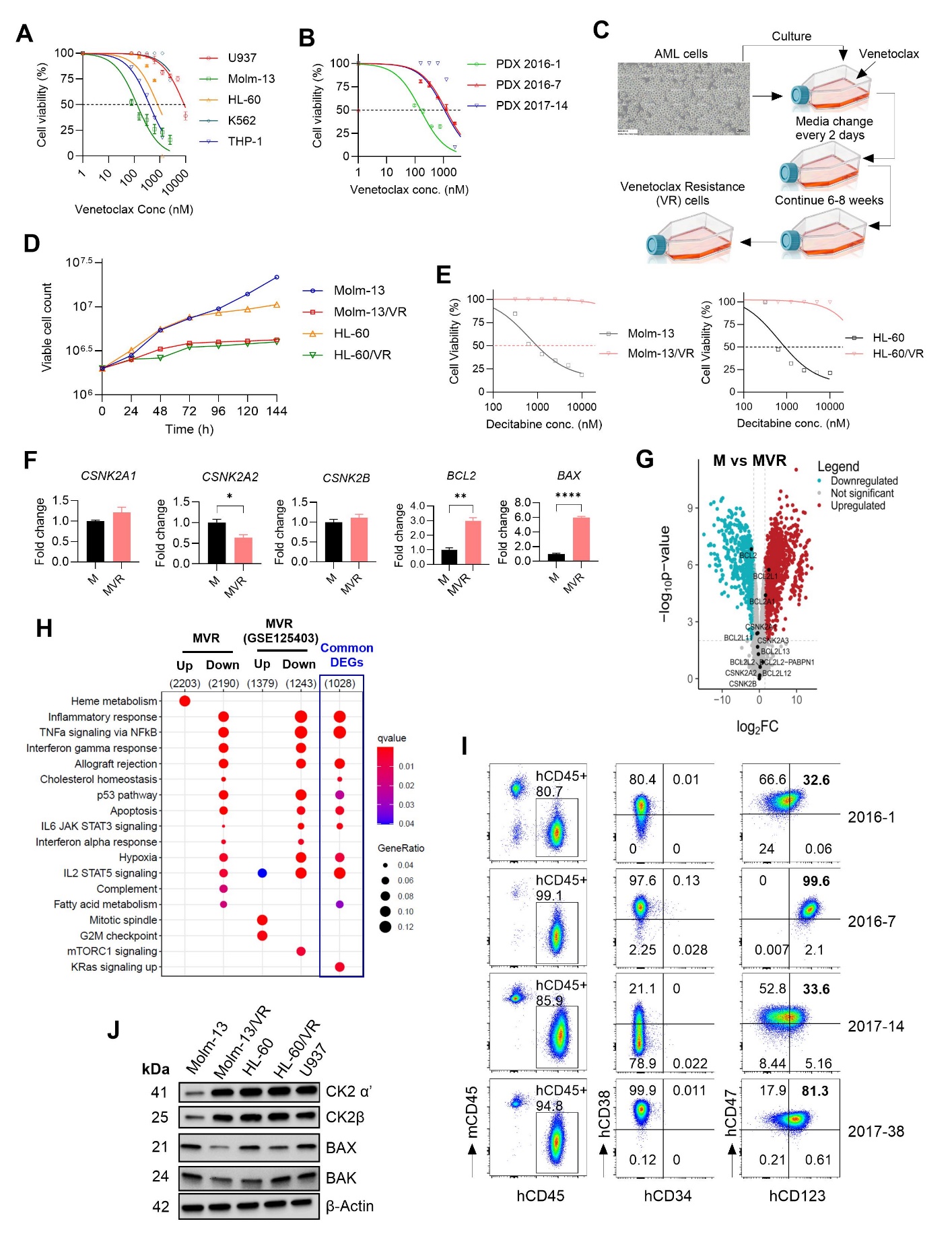

**Supplementary Figure 1: The characterization of venetoclax-resistant AML cell lines and cytotoxic effect of different standard drugs on AML cell lines and PDX cells. A-B)** AML cell lines (A) and PDX cells (B) were treated with different concentrations of venetoclax for 48 h and accessed for cell viability using WST assay. **C)** Schematic flow chart describing the generation of VEN-resistant cell lines from parental VEN-sensitive AML cells. **D)** The proliferation of venetoclax susceptible (parental) and VEN-resistant (VR) AML cells was analyzed for 6 days using CellDrop automated cell counter. **E)** The cross resistance of venetoclax-resistant AML cells was tested against different concentrations of decitabine for 48 h and assessed for cell viability. **F)** The basal level expression of indicated genes was assessed in different AML cell lines by qRT-PCR. The data are presented as mean ± SEM (n=3 tech replicates from a representative run). *p<0.05, **p<0.01 and ****p<0.0001 by unpaired t-test (Welch’s correction) denotes statistical significance. **G)** Differential gene expression analysis by RNA-seq in Molm-13/VR (MVR) compared to parental Molm-13 (M) cell line was shown in the volcano plot. **H)** Dot plot showing the top-ranked functional pathways (MSigDB hallmark gene set) enriched in differentially regulated genes (fold-change >1.5 and adj-p <0.05) in MVR cells when compared to parental cells. The molecular gene signatures in our MVR cells are similar and comparable to that of MVR cells transcriptome from a different study (GSE125403). **I)** The basal level surface expression of markers for LSCs (CD34, CD38) and chemoresistance (CD47, CD123) was analyzed using flow cytometry in different AML PDX cells. **J)** The basal level expression of indicated target proteins in different AML cell lines was assessed by immunoblotting.

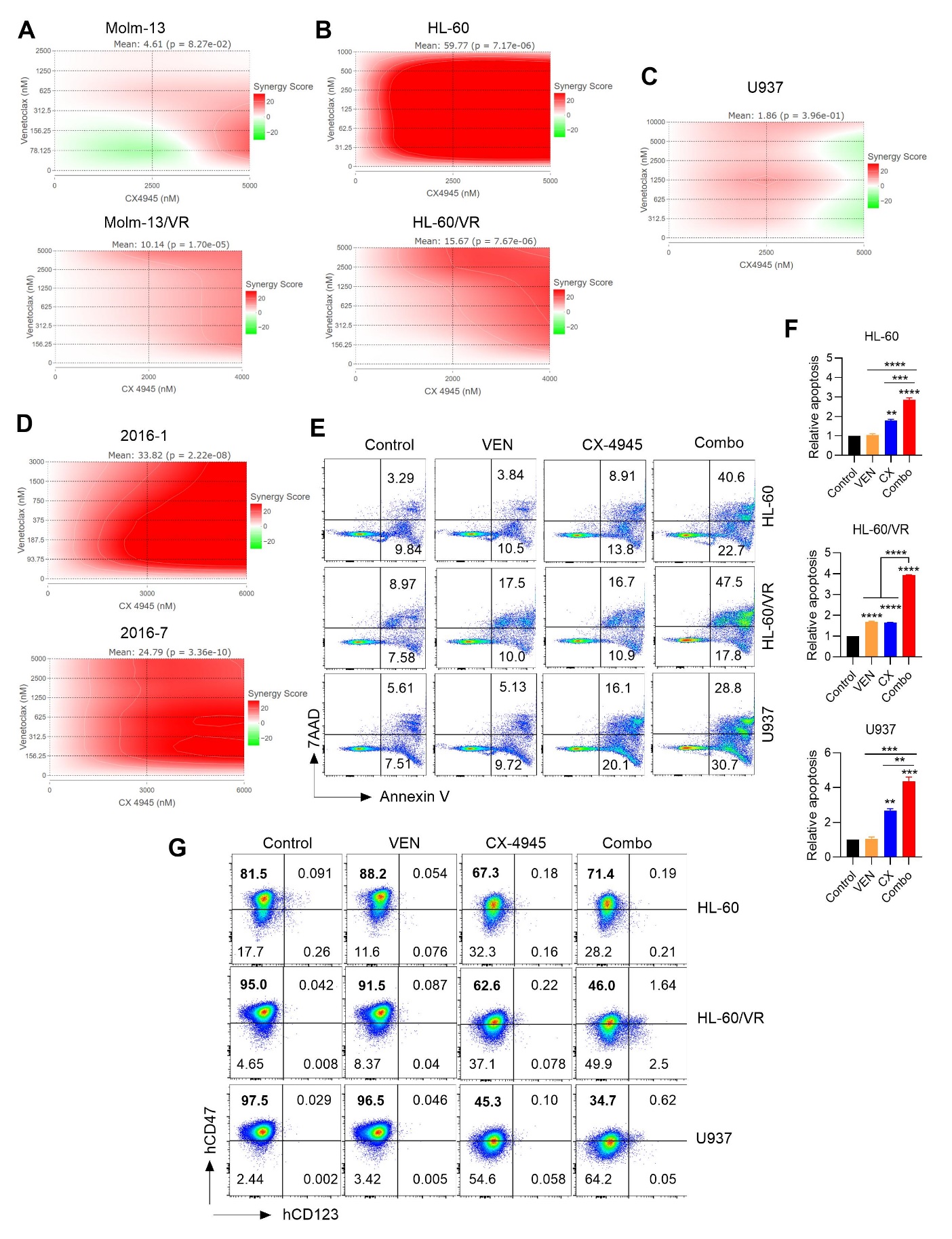

**Supplementary Figure 2: The synergistic cytotoxic effect of CX-4945 in combination with VEN in parental and VR-AML cell lines. A-D)** A representative synergy map showing the cytotoxic effect of CX-4945 and VEN combination in the AML cell lines Molm-13 parental and Molm-13/VR (**A**), HL-60 parental and HL-60/VR (**B**), U937 (**C**), 2016-1 and 2016-7 PDX cells (**D**) cells treated for 48 h. were treated with different concentrations of VEN in combination with CX-4945 for 48 h and cell viabilities were assessed by WST assay. ZIP Synergy scores were calculated using the SynergyFinder Plus (<https://synergyfinder.org/>) interactive webtool. ZIP scores greater than 10 indicates ‘synergy’, and 0-10 indicate ‘additive’ activity between CX-4945 and VEN. **E-F)** AML cell lines (HL-60, HL-60/VR, U937) were treated with CX-4945 and VEN alone or in combination for 24 h and stained with Annexin V/7AAD for analyzing apoptosis by flow cytometry (**E**). Relative apoptosis was calculated by normalizing to vehicle treated cells (**F**). The data are presented as mean ± SD (n=2). **p<0.01, ***p<0.001 and ****p<0.0001 by one-way ANOVA (Tukey’s multiple comparisons test) denotes statistical significance. **G)** AML cell lines (HL-60, HL-60/VR, U937) were treated with CX-4945 and VEN alone or in combination for 24 h and the surface expression of chemoresistance markers (CD47 and CD123) was analyzed by flow cytometry.

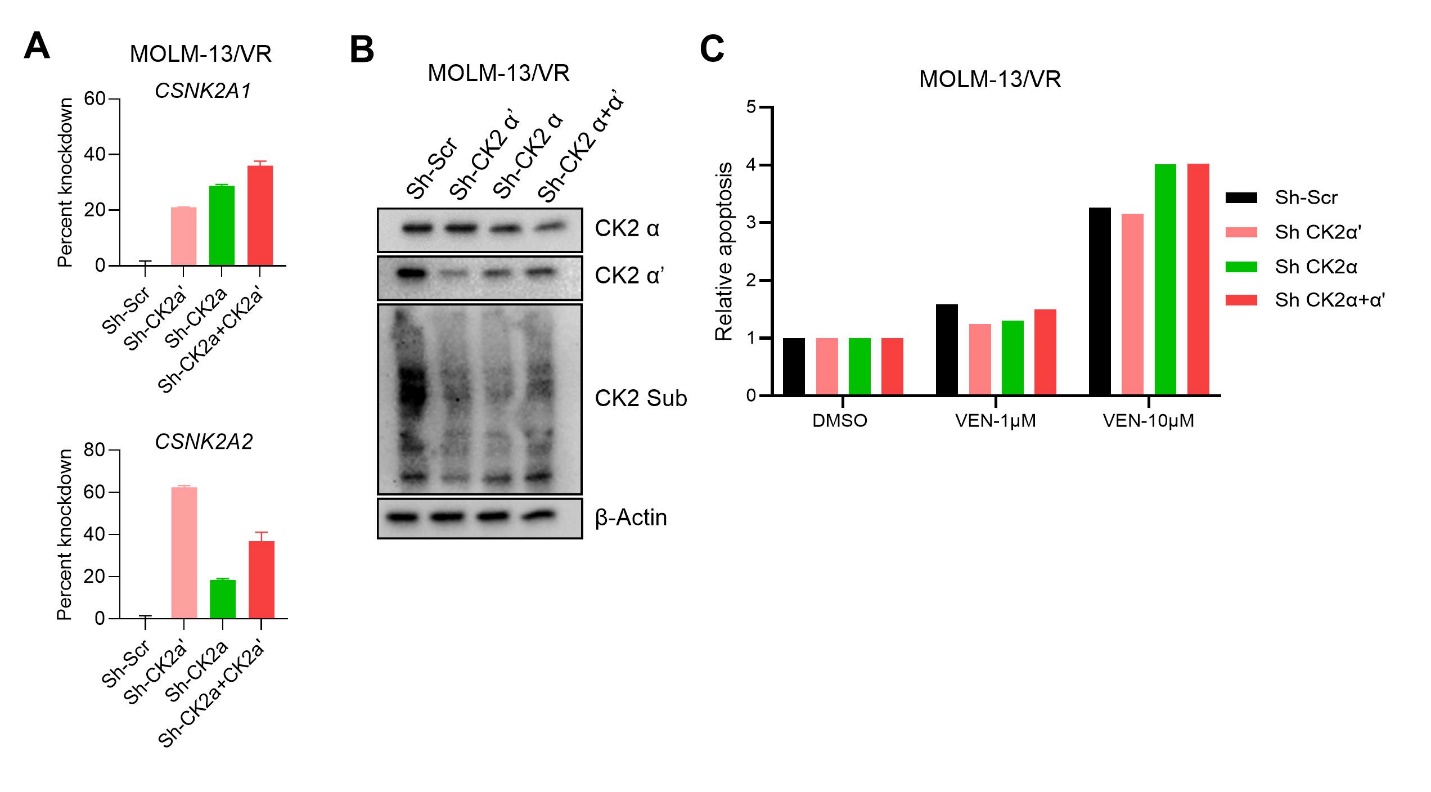

**Supplementary Figure 3: The effect of shRNA-mediated knockdown of *CSNKA1* and *CSNKA2* genes on the activity of VEN in VR-AML cells. A-B)** Molm-13/VR cells were transduced with lentiviral particles expressing GFP-tagged shRNAs specific to *CSNK2A1* and *CSNK2A2* alone or together in the presence of polybrene (8 µg/mL). The cells positive for GFP were sorted and tested for target gene knockdown using qRT-PCR (**A**) and western blotting (**B**). β-Actin was used as a loading control for western blotting experiments. **C)** The relative apoptosis in Molm-13/VR cells after knockdown of *CSNK2A1* and *CSNK2A2* genes alone or combination was analyzed after 24 h treatment with indicated doses of VEN using muse cell analyzer.

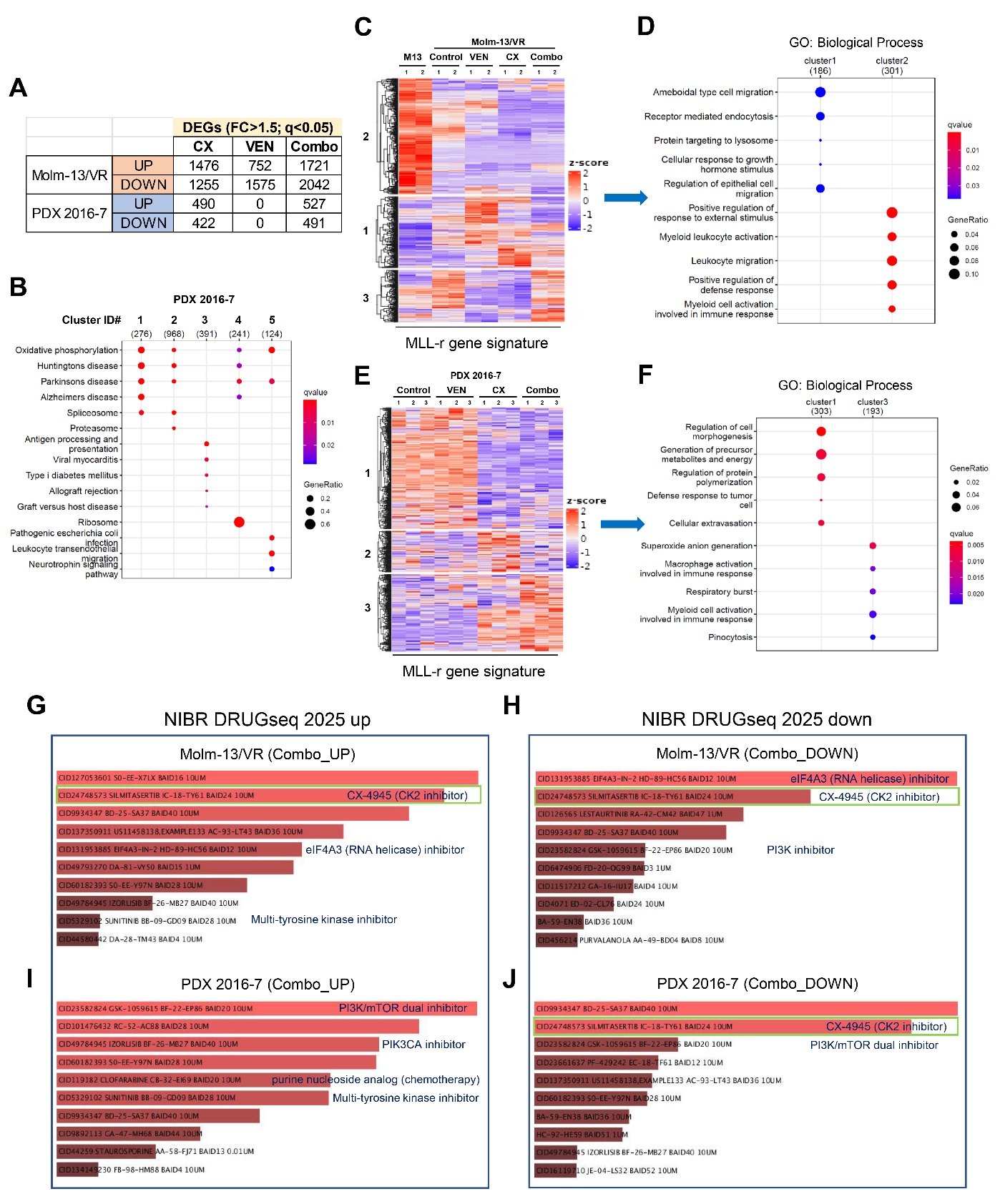

**Supplementary Figure 4: Enrichment analyses of genes differentially regulated in Molm-13/VR and PDX 2016-7 cells after treatment with CX-4945, VEN, and Combo. A)** Summary plot showing the number of differentially expressed genes (DEGs) in Molm-13/VR and PDX 2016-7 cells after 24 h of treatment with CX-4945 (CX), venetoclax (VEN), and CX+VEN combination (Combo). **B)** Dot plot showing the top-ranked biological processes (gene ontology) enriched in different gene clusters obtained with k-means clustering of PDX 2016-7 cells transcriptome (top 2000 most-variable genes). **C-F)** k-means clustering approach separated 623 MLL-r signature genes (PMID: 17597811 & 4D PDX source paper) based on their expression in Molm-13 and Molm-13/VR (**C**), and PDX 2016-7 cells (**E**). Subsequent enrichment analysis of different gene clusters obtained with k-means clustering of MLL-r genes showed top-ranked biological processes (gene ontology) in bubble plot (**D, F**). **G-J)** DrugSeq (Drug perturbation signatures) enrichment analysis bar plot of DEGs obtained by Enrichr web tool. The genes upregulated (**G, I**) and downregulated (**H, J**) after treatment with CX+VEN combination in Molm-13/VR (**G, H**) and 2016-7 PDX cells (**I, J**) were checked against most recent NIBR_DRUGseq_2025 gene-set libraries that significantly overlaps with input genes. The length of the bar and intensity of the color represents the significance of that specific gene-set or term.

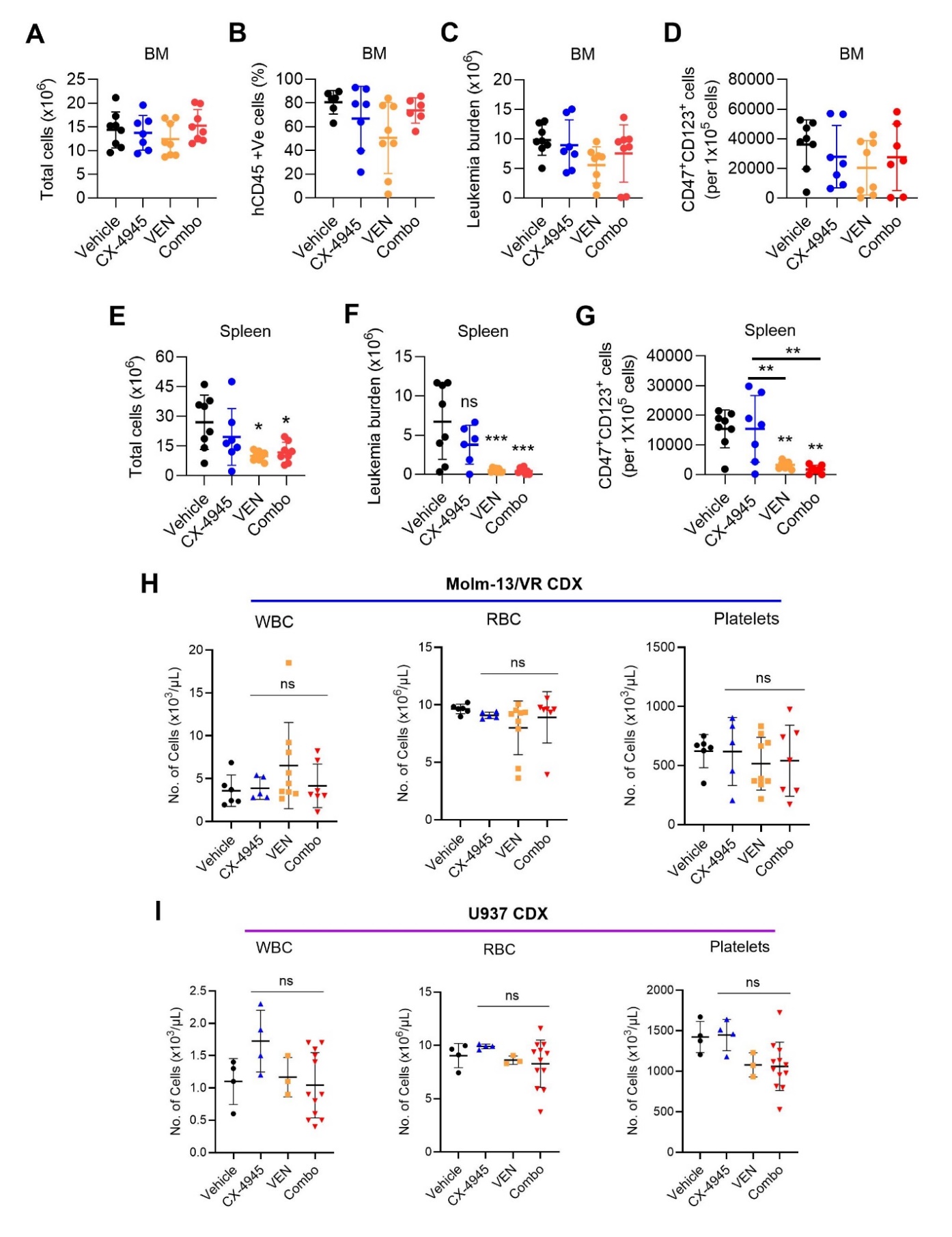

**Supplementary Figure 5: The effect of CX-4945 and VEN combination on complete blood count and leukemia burden in mice xenografted with Molm-13/VR, U937, and PDX 2016-7 cells. A-G)** NRG-S mice transplanted with AML PDX 2016-7 cells (0.5x10^6^ cells/mouse) intravenously were treated with CX-4945, VEN, and Combo. The tissues (spleen, bone marrow) were collected from PDX mice after two-weeks of drug treatment and subjected to flow cytometric analysis of total cell number (**A, E**), leukemia burden (**B, C, F**), and chemoresistant cells (**D, G**). The data are presented as mean ± SD (n=7-8). **H & I)** NRG-S mice transplanted with Molm-13/VR (0.25x10^6^ cells/mouse) **(G)** and U937 (1x10^4^ cells/mouse) (**H**) cells intravenously were treated with CX-4945, VEN, and Combo. After two-weeks of drug treatment, complete blood count (CBC) was analyzed from Molm-13/VR (**G**) and U937 (**H**) CDX mice by Hemavet analyzer. Data are presented as mean ± SD (n=4-12). *p<0.05, **p<0.01, and ***p<0.001 by one-way ANOVA (Tukey’s multiple comparisons test) denotes statistical significance.

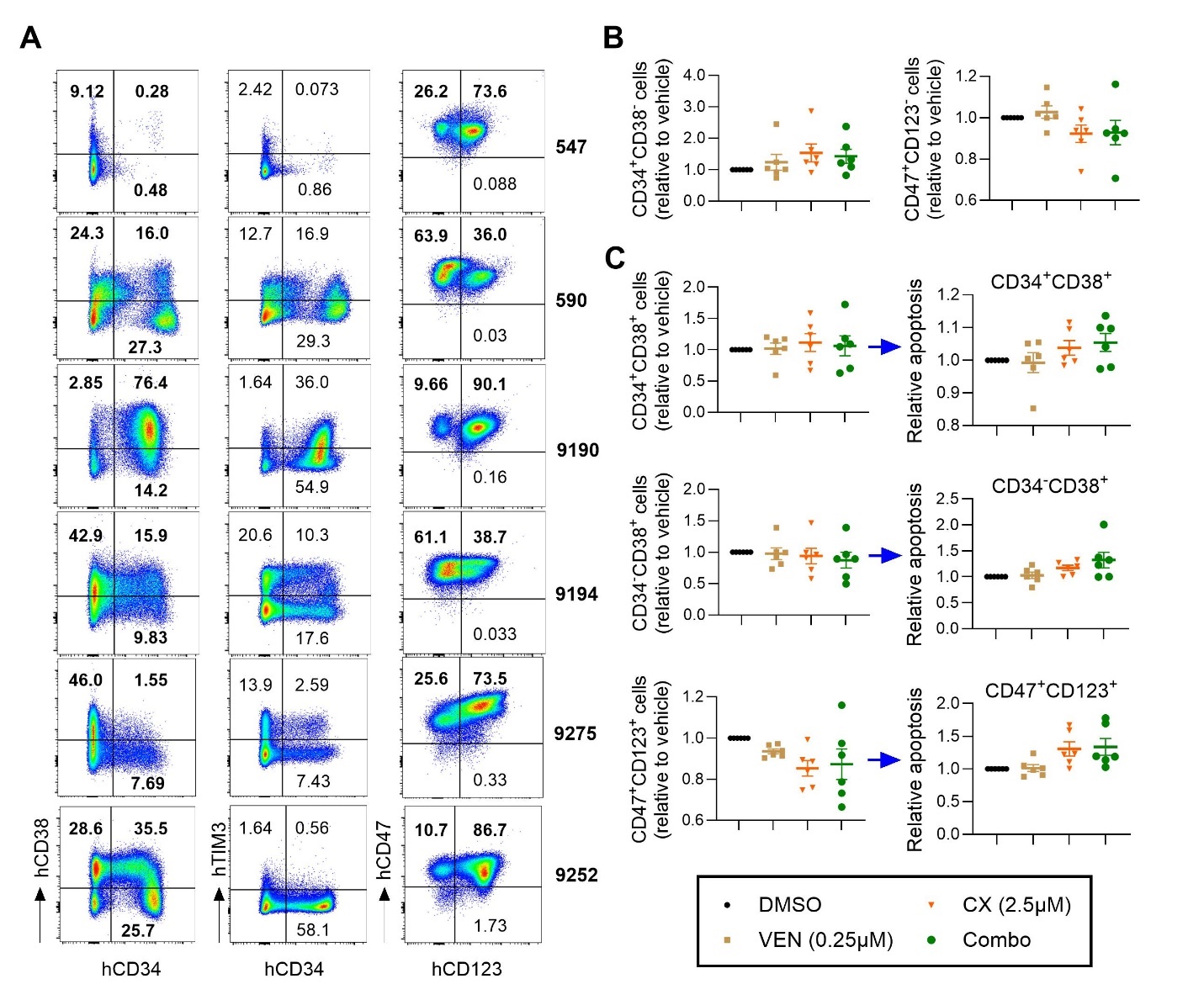

**Supplementary Figure 6: Characterization and testing the effect of CX-4945 and VEN combination in AML patient cells. A)** AML cells from patients were surfaced stained with different fluorescent labeled antibodies and analyzed for the expression of markers for LSCs (CD34, CD38, TIM3) and chemoresistance (CD47 and CD123) by flow cytometry. **B & C)** AML patient cells were treated with CX-4945 in combination with VEN for 24 h and surface stained for different cell surface markers along with annexin V. The relative cell numbers (**B & C**) and apoptotic cells (**C**) were calculated by normalizing to vehicle treated cells. The data are presented as mean ± SEM (n=6) and analyzed via one-way ANOVA (Tukey’s multiple comparisons test). *p<0.05 considered as statistically significant.

**Table S1: Characteristics of AML cell lines**

| **Cell line** | **Source**  **(Cat. No.)** | **Tissue** | **Disease** | **Age** | **Gender** | **STR DNA Profile** |
| --- | --- | --- | --- | --- | --- | --- |
| **U937** | ATCC (CRL1593.2) | Pleural effusion, lymphocyte, myeloid | Histiocytic lymphoma | 37 yrs | Male | Amelogenin: X; CSF1PO: 12; D13S317: 10,12; D16S539: 12; D5S818: 12; D7S820: 9,11; TH01: 6, 9.3; TPOX: 8,11; vWA: 14, 15 |
| **Molm-13** | DSMZ  (ACC 554) | Peripheral blood | Acute myeloid leukemia | 20 yrs | Male |  |
| **THP-1** | ATCC  (TIB-202) | Peripheral blood | Acute monocytic leukemia | 1 yrs | Male | Amelogenin: X,Y; CSF1PO: 11,13; D13S317: 13, D16S539: 11,12; D5S818: 11,12; D7S820: 10; TH01: 8,9.3; TPOX: 8,11; vWA: 16 |
| **HL-60** | ATCC  (CCL-240) | Peripheral blood | Acute promyelocytic leukemia | 36 yrs | Female | D5S818: 12; D13S317: 8,11; D7S820: 11,12; D16S539: 11; vWA: 16; TH01: 7,8; Amelogenin: X; TPOX: 8,11; CSF1PO: 13,14 |
| **K562** | ATCC (CCL243) | Bone marrow | Chronic myelogenous leukemia | 53 yrs | Female | Amelogenin: X; CSF1PO: 9,10; D13S317: 8; D16S539: 11,12; D5S818: 11,12; D7S820: 9,11; TH01: 9.3; TPOX: 8,9; vWA: 16 |

**Table S2: Characteristics of AML Primary samples**

| **Group** | **Sample ID** | **Diagnosis** | **Age** | **Gender** | **Cytogenetics** |
| --- | --- | --- | --- | --- | --- |
| MLL | 9252 | AML | 32 | M | 46,XY,t(6;11)(q27;q23)[20] |
| FLT3-ITD | 591 | AML | 62 | F | 46,XX[20] FLT3-ITD, IDH1, KDM6A, NPM1, and DNMT3A |
| FLT3-ITD | 674 | AML | 58 | M | 46,XY[20] FLT3-ITD, DNMT3A, and SMC3 mutated |
| FLT3-ITD | 609 | AML | 50 | F | 46,XX[20] DNMT3A, TET2, NPM1, FLT3-ITD |
| FLT3-ITD | 686 | AML | 61 | F | 46,XX,t(6;9)(p23;q34)[20] FLT3-ITD |
| FLT3-ITD | 494 | AML | 60 | M | 46,XY[20] FLT3-ITD, NPM1 mutations |
| Normal cyto | 771 | AML | 26 | F | 46,XX[20] |
| Normal cyto | 9192 | MDS/AML | 64 | F | 46,XX[20] |
| Normal cyto | 9215 | AML | 62 | M | 46,XY[20] |
|  | 547 | MDS/AML | 65 | M | 46,XY,del(20)(q11.2q13.3)[5]/46,XY[15]  IDH2, NPM1, and SRSF2 mutations |
| Complex | 590 | AML | 71 | F | 45-50,XX,del(5)(q15q33),t(6;20)(p21.3;q13.1),-7,+add(8)(p11.2),add(8)(p21),dic(8;21)(p21;q22),i(8)(q10),-18,+dic(21;21)(q22;q22)x1-2,+0-2r,+0-1mar[cp19]/46,XX[1] TP53 |
| FLT3-ITD | 9190 | AML | 46 | F | 47,XX,add(1)(p36.1),t(6;9)(p23;q34),add(12)(p13),+15[20] FLT3-ITD (71.6%) |
|  | 9194 | AML | 45 | M | 47,XY,inv(16)(p13.1q22),+22[20] |
| Normal cyto | 9275 | AML with monocytic phenotype | 68 | F | 46,XX[20] |
|  | 720 | AML | 52 | F | 46,XX,add(11)(p15)[4]/46,idem,add(2)(q31),del(11)(q23),i(17)(q10)[16] DNMT3A, FLT3-TKD |
| Complex | 9296 | AML | 30 | M | 43-45,XY,add(3)(q13.2),-5,der(5;13)(p10;q10),-7,-12,-20,-20, add(19)(q13.3),+3-6mar[cp20] |
| Normal cyto | 9351 | AML | 32 | M | 46,XY[20] |

Patient cells listed in shaded area (n=7) were used for the study. Other patient cells were omitted from study due to high basal level apoptosis and poor viability post-thaw.

**Table S3: Characteristics of PDX AML cells**

| **Patient ID** | **PDX Model** | **Diagnosis** | **Age/Sex** | **MLL Status** | **Genetics** | **Source Reference** |
| --- | --- | --- | --- | --- | --- | --- |
| 2017-38 | AML-09 | R/R AML | Unknown | MLL-r | MLL/AF9, NRAS | (Wunderlich et al., 2022) |
| 2016-1 | AML-10 | R/R AML | 14 y /F | Wildtype | CALM-AF10 | (Wunderlich et al., 2022) |
| 2016-7 | AML-18 | R/R AML | 16y /F | MLL-r | MLL/AF9 | (Wunderlich et al., 2022) |
| 2017-14 | AML-19 | R/R tAML | 6y /F | MLL-r | MLL/AF10 | (Wunderlich et al., 2022) |

**Table S4: Primers used for qRT-PCR**

| **Target** | **Forward primer (5’ 🡪 3’)** | **Reverse primer (5’ 🡪 3’)** | **Source** |
| --- | --- | --- | --- |
| CK2α  (*CSNK2A1*) | GGTGAGGATAGCCAAGGTTCTG | TCACTGTGGACAAAGCGTTCCC | IDT |
| CK2α’  (*CSNK2A2*) | CGACCATCAACAGAGACTGACTG | GTGAGACCACTGGAAAGCACAG | IDT |
| CK2b (*CSNK2B*) | CAGAGTGACCTGATTGAGCAGG | CGAGGACAGTAACCAAAGTCTCC | IDT |
| *BCL2* | ATCGCCCTGTGGATGACTGAGT | GCCAGGAGAAATCAAACAGAGGC | IDT |
| BCL-XL (*BCL2L1*) | GCCACTTACCTGAATGACCACC | AACCAGCGGTTGAAGCGTTCCT | IDT |
| *MCL1* | CCAAGAAAGCTGCATCGAACCAT | CAGCACATTCCTGATGCCACCT | IDT |
| *BAK* | TTACCGCCATCAGCAGGAACAG | GGAACTCTGAGTCATAGCGTCG | IDT |
| *BAX* | TCAGGATGCGTCCACCAAGAAG | TGTGTCCACGGCGGCAATCATC | IDT |
| *BCL2A1* | GGATAAGGCAAAACGGAGGCTG | CAGTATTGCTTCAGGAGAGATAGC | IDT |
| *GAPDH* | GTCTCCTCTGACTTCAACAGCG | ACCACCCTGTTGCTGTAGCCAA | IDT |

**Table S5: List of antibodies used for flow cytometry analysis of AML cells**

| **Target** | **Fluorochrome** | **Clone** | **Reference** | **Species** | **Provider** |
| --- | --- | --- | --- | --- | --- |
| mCD45 | BV711 | 30-F11 | 103147 | Rat IgG2b, κ | BioLegend |
| hCD47 | APC | B6H12 | 17-0479-42 | Mouse IgG1, κ | eBioscience |
| hCD123 | PE/Cy7 | 6H6 | 983702 | Mouse IgG1, κ | BioLegend |
| hCD45 | FITC | HI30 | 304006 | Mouse IgG1, κ | BioLegend |
| hCD34 | PE | 581 | 343506 | Mouse IgG1, κ | BioLegend |
| hCD38 | APC/Cy7 | HB-7 | 356616 | Mouse IgG1, κ | BioLegend |
| hCD34 | FITC | 581 | 343504 | Mouse IgG1, κ | BioLegend |
| hCD366 (Tim-3) | BV421 | A18087E | 364808 | Mouse IgG1, κ | BioLegend |

**Table S6: List of antibodies used for immunoblotting analysis**

| **Target** | **Clone** | **Reference** | **Provider** |
| --- | --- | --- | --- |
| CK2α | E-7 | SC-373894 | Santa Cruz |
| p-CK2 substrate |  | 87385 | Cell Signaling |
| p-AKT (S129) | D4P7F | 13461s | Cell Signaling |
| p-NFκB p65 (S536) | 93H1 | 3033 | Cell Signaling |
| PARP |  | 9542s | Cell Signaling |
| BCL-XL | 54H6 | 2764s | Cell Signaling |
| MCL-1 (detects L, S, ES isoforms) | D35A5 | 5453s | Cell Signaling |
| MCL-1 (detects only L isoform) | D5V5L | 39224 | Cell Signaling |
| BCL2 | C-2 | SC-7382 | Santa Cruz |
| BCL2A1 (A1/Bfl-1) | E3U2F | 24951s | Cell Signaling |
| BAK | D2E11 | 5023s | Cell Signaling |
| BAX | D4E4 | 12105s | Cell Signaling |
| β-actin | 13ES | 4970s | Cell Signaling |
| GAPDH | 14C10 | 2118 | Cell Signaling |
